# π-MSNet: A large-scale, AI-ready and continuously reprocessed proteomics data resource

**DOI:** 10.64898/2026.04.13.718149

**Authors:** Chengxin Dai, Yi Liu, Tianze Ling, Qingyuan Zhang, Shouyang Ren, Yuan Chu, Menglu Wang, Yang Qiu, Huali Xu, Xiaowei Huang, Yunping Zhu, Timo Sachsenberg, Mingze Bai, Fuchu He, Yasset Perez-Riverol, Linhai Xie, Cheng Chang

## Abstract

Artificial intelligence (AI) is reshaping proteomics workflows, delivering remarkable gains in both peptide identification sensitivity and quantitative performance. However, the potential of deep learning models in proteomics has not been fully exploited due to the scarcity of large-scale, high-quality and consistently labeled datasets. Here, we present π-MSNet, a large-scale, AI-ready and continuously reprocessed mass spectrometry (MS) data resource. Using a uniform identification and quality control workflow, it comprises over 737 million peptide-spectrum matches (PSMs) obtained from 2.16 billion MS2 spectra, and 11 million precursors from 45,583 LC-MS/MS runs across 12 instrument types and 55 diverse species. Through community collaboration, the data are shared via international, interactive, and continuously reprocessed web resources. π-MSNet provides an AI-ready data framework for efficient training and systematic benchmarking of multiple models across four representative tasks (e.g., MS2 prediction, retention time prediction, collisional cross sections prediction, and de novo peptide sequencing). In particular, by retraining multiple models on π-MSNet, we achieved consistent performance improvements over their original versions. These improved models were subsequently integrated into the π-MSNet agent to enable interactive, deployment-free use. Through SDRF (Sample and Data Relationship Format) metadata, open-source cloud analysis workflow, and a community-driven interactive data portal that supports continuous data submission, π-MSNet serves as a continuously updating, AI-ready resource for reproducible benchmarking, robust model training, and accelerated AI innovation in proteomics.

## Main

In the rapidly evolving field of computational proteomics, data-driven methodologies, particularly deep learning, have become increasingly indispensable for interpreting increasingly complex mass spectrometry (MS) data^1^. Over the past decade, the latest neural network architectures have been integrated into nearly every stage of the proteomics data analysis pipeline, fundamentally reshaping how peptide and protein identifications are performed. For example, deep neural networks are now widely used to predict fragment ion intensities and liquid chromatography (LC) retention time (RT), which are subsequently incorporated into peptide-spectrum match (PSM) rescoring frameworks to further improve identification confidence and proteome coverage^2–4^. In parallel, deep learning has enabled the development of powerful de novo peptide sequencing algorithms that infer peptide sequences directly from tandem MS spectra without reliance on sequence databases^5,6^. Collectively, these advances demonstrate the transformative potential of artificial intelligence (AI) in proteomics.

Despite the wealth of publicly accessible raw MS data, which are largely centralized within the ProteomeXchange consortium^7^—with PRIDE^8^ and iProX^9^ serving as key contributors to the growth and accessibility of proteomics datasets—current repositories primarily store raw files with heterogeneous and incomplete metadata annotations and inconsistent processing standards. These limitations complicate the direct use of such datasets in machine learning workflows. Consequently, there remains a critical shortage of large-scale, high-quality, diverse, and AI-ready datasets specifically tailored for deep learning model training and benchmarking. An AI-ready dataset in proteomics is a curated, standardized, and computationally accessible collection of mass spectrometry–derived data and associated metadata that adheres to the FAIR (Findable, Accessible, Interoperable, and Reusable) principles^10^, and is explicitly structured to enable efficient, reproducible, and scalable machine learning. Moreover, the absence of standardized evaluation datasets hinders fair comparison across computational methods and undermines reproducibility^11^. This fragmentation exacerbates the opacity of AI models, limits their generalizability across experimental settings, and ultimately constrains their widespread and large-scale adoption in proteomics research^12,13^.

Although several annotated PSM resources have been established, their scale and diversity remain insufficient to reflect the complexity of contemporary proteomics experiments. For instance, MassIVE-KB^14^, released in 2016 and currently the largest publicly available annotated PSM dataset, contains approximately 30 million PSMs and 2 million precursor ions; however, it is restricted to higher-energy collisional dissociation (HCD) spectra acquired on Orbitrap instruments and predominantly includes peptides with carbamidomethylated cysteines as the primary chemical modification. Such constraints limit its representativeness with respect to diverse fragmentation methods, instrument platforms, sample preparation strategies, and post-translational modifications (PTMs). Furthermore, MS data generated from emerging experimental paradigms—such as single-cell proteomics and trapped ion mobility spectrometry (e.g. timsTOF)—are increasingly deposited in ProteomeXchange repositories. These evolving modalities introduce new spectral characteristics, noise profiles, and analytical challenges that are not captured by existing static datasets. Consequently, there is a pressing need for a continuously updated, standardized, and scalable resource capable of accommodating the expanding diversity of MS data.

Here, we present π-MSNet, a standardized, large-scale, AI-ready, and continuously reprocessed dataset resource designed to accelerate the development, training, and benchmarking of computational methods in proteomics. We curated and annotated 127 large-scale proteomics datasets from ProteomeXchange and the π-HuB project^15^ in SDRF^16^ format, encompassing diverse biological contexts, instrument platforms, and fragmentation strategies. All raw MS datasets were systematically and continuously reanalyzed using quantms^17^ for DDA data and quantmsdiann for DIA data^18^, both of which are open-source, cloud-enabled, and fully reproducible workflows for identification and quantification. The resulting annotations were harmonized and stored in Quantitative Proteomics eXchange (QPX), a unified, structured Parquet format optimized for scalable machine learning applications. Beyond data integration, we retrained and benchmarked existing state-of-the-art models on the expanded, diversified PSM collection to demonstrate performance improvements achieved with π-MSNet across downstream tasks. π-MSNet is distributed via interactive web portals at https://msnet.ncpsb.org.cn and https://portal.quantms.org, and a companion AI agent powered by retrained and calibrated models is publicly available at https://msnet.ncpsb.org.cn to facilitate user-friendly analysis of MS datasets. Together, π-MSNet provides a scalable open resource that supports the development of AI methods and community-standard benchmarking in computational proteomics.

## Results

Here, we introduce π-MSNet, a large-scale, AI-ready and continuously reprocessed data resource for proteomics (**Figure 1a**). The π-MSNet resource was compiled from public datasets available at the ProteomeXchange consortium data repositories including iProX, PRIDE Archive and the in-house data from the π-HuB project. Collectively, these datasets comprise over 2.16 billion tandem mass spectrometry (MS2) spectra from 45,583 LC-MS/MS runs across 127 datasets, with a total size of 40 terabytes (TB). Prior to analysis, all collected datasets were uniformly annotated in SDRF format, the HUPO-PSI standard metadata format.

**Fig. 1.**
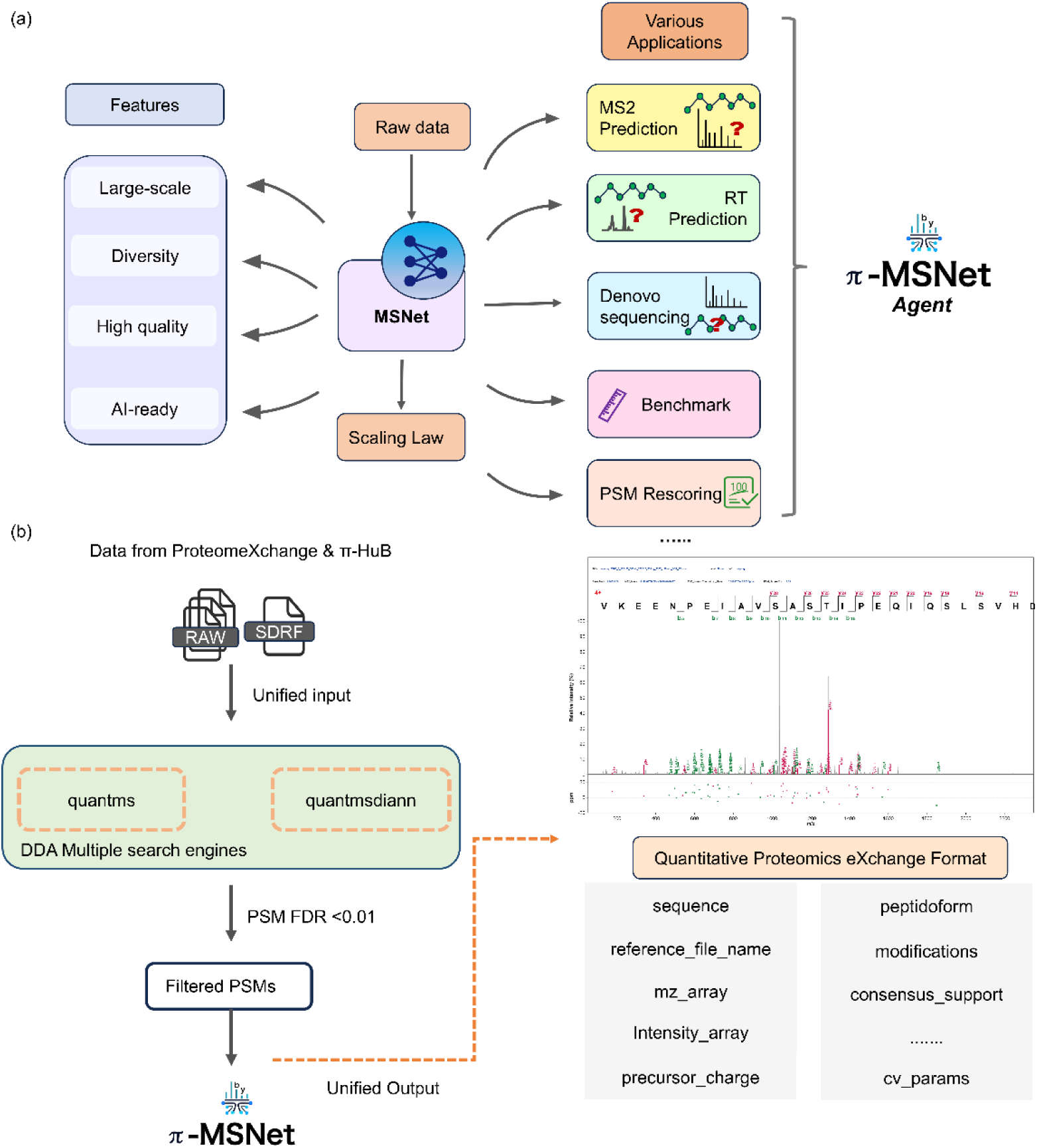
| **π-MSNet as a foundational resource for proteomics downstream tasks. (a)** π-MSNet can then be adapted to a wide range of downstream tasks in proteomics, all of which are incorporated into the π-MSNet agent. **(b)** π- MSNet processing workflow: reanalysis workflow accepts SDRF metadata files, raw mass spectrometry data, and FASTA-formatted protein sequence databases as input. All DDA MS datasets were reanalyzed using multiple search engines in combination with Percolator^21^ to mitigate engine-specific biases, and subsequently filtered at a PSM false discovery rate (FDR) of 1%. The DIA dataset was analyzed using quantmsdiann, configured with DIA-NN v2.5.1^22^. For the PTM dataset, an additional false localization rate (FLR) threshold of <0.01 was applied.

For DDA datasets, we reanalyzed the MS2 spectra using the quantms workflow. By integrating PSMs from multiple search engines (e.g., MS-GF+^19^ and Comet^20^) into a consensus score for each PSM, we mitigated engine-specific biases and enhanced the robustness of peptide-spectrum annotations. For DIA datasets, we reanalyzed the MS2 spectra using the quantmsdiann workflow. These efficient and standardized data-processing pipelines underpin the scalability of π-MSNet, enabling large-scale reanalysis and release of public proteomics datasets. Across representative projects spanning a wide range of data volumes, the pipelines supported end-to-end processing and release at different scales. For example, the end-to-end reanalysis and release of 58,527,901 PSMs was completed within 26h using Kubernetes cluster with six nodes.

The generated PSM results were exported in the unified space-efficient QPX Parquet format for rapid data access and exchange (https://github.com/bigbio/qpx), where each PSM entry consists of PSM annotation columns and metadata columns. Among them, the metadata columns primarily store values about the instrument, collision method, collision energy, and mass analyzer (**Figure 1b**). Compared to CSV and HDF5, the QPX format reduces storage space by 75% and 96%, and reading times by 50% and 90% respectively.

Finally, we established the π-MSNet data portal, which includes 737 million PSMs and 11 million precursors from 55 species, including eukaryotes, prokaryotes, viruses, and archaea (**Figure 2**). These spectral data were acquired on 12 different types of mass spectrometers, yielding a wide and diverse data resource from real experiments. In addition to 709 million PSMs from typical tryptic peptides, π-MSNet contains 24.9 million PSMs from non-specific cleavage peptides, 1.47 million PSMs from Lys-C digestion, 503,534 PSMs from Glu-C digestion, and 148,896 PSMs from chymotrypsin digestion. Therefore, the π-MSNet covers a broader range of peptide space: peptide lengths from 6 to 40 and precursor charges from 1 to 6.

**Fig. 2.**
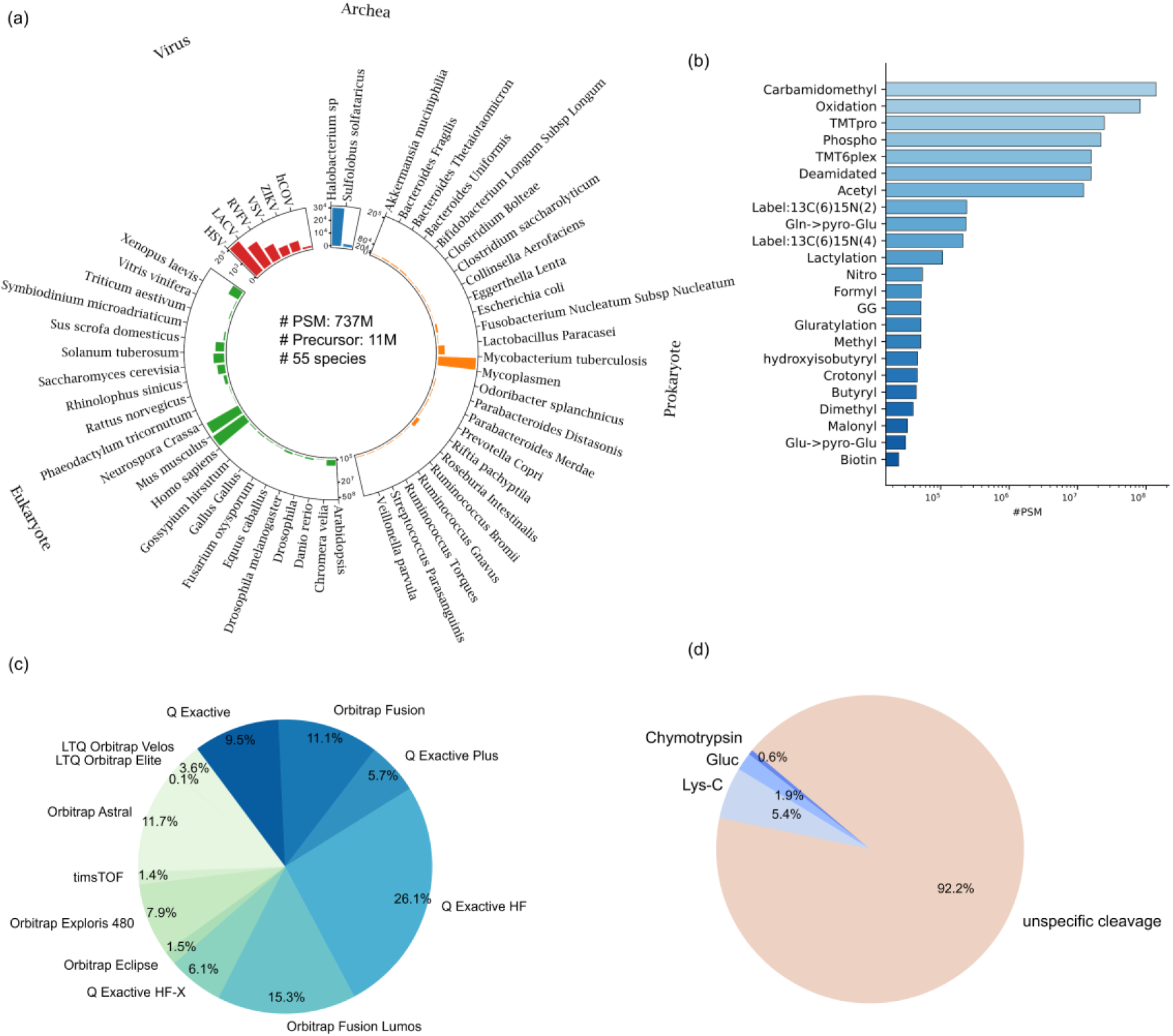
| **π-MSNet Overview**. (a) The number of PSM from different species, including eukaryotes (green), virus (red), archaea (blue) and prokaryotes (orange). (b) The number of PSMs for 23 modification types. (c) Distribution of PSMs across 12 instrument types, ordered clockwise by market introduction time. (d) Distribution of non-tryptic PSMs across four cleavage methods, highlighting the distribution of non-tryptic peptides.

To address the diverse data and feature requirements of downstream proteomics tasks, we developed MSNetLoader (https://github.com/PHOENIXcenter/pi-MSnet), a Python library that enables seamless, flexible integration of π-MSNet data into AI models, thereby supporting AI-ready usage. The continuously reprocessing architecture of π-MSNet enables continuous expansion to keep pace with the rapid growth and diversification of public MS data. Instead of relying on static snapshots, π- MSNet supports incremental updates through unified, cloud-based reanalysis workflow built on quantms and quantmsdiann with standardized SDRF metadata, ensuring consistent processing and harmonized annotations across newly integrated datasets. Through continuous reprocessing of newly incorporated datasets, π-MSNet has further expanded in both data volume and experimental diversity, extending beyond LFQ DDA datasets to encompass DIA, TMT- and SILAC-labeled, and metaproteomics datasets. The same standardized workflows can also be adopted by the community to generate and submit QPX-formatted results, providing a route for community-contributed datasets to be incorporated into π-MSNet and further supporting its continued expansion. Together, these mechanisms establish π-MSNet as a scalable, extensible, and continuously reprocessed resource for reliable large-scale machine learning training and evaluation in proteomics.

We further demonstrated the utility of π-MSNet as an AI-ready data portal by training and benchmarking models on four representative proteomics tasks: fragmentation pattern prediction (MS2 prediction), chromatographic behavior modeling (retention time prediction), collisional cross sections prediction (CCS prediction) and direct sequence inference (de novo peptide sequencing).

At the signal level, MS2 prediction models the fragmentation patterns of peptides. In MS2 prediction tasks, we began by verifying scaling laws based on AlphaPeptDeep architecture and observed that the test loss decreases continuously as the numbers of PSMs and precursors in the training data increase synchronously (**Figures 3a-c**). We benchmarked three popular models, i.e. AlphaPeptDeep^2^, Prosit^23^ and Unispec^24^, and considered only b+, b++, y+, and y++ ions, as common methods mainly focus on accurately predicting the fragment intensities of b and y ions. AlphaPeptDeep achieved a higher proportion of Pearson correlation coefficients above 0.9 (PCC90) than the other two models across multiple test datasets (**Figures 3d-e**). More importantly, retraining AlphaPeptDeep with progressively scaled π-MSNet training data improved its average PCC90 from 0.68 to 0.85 compared to the published version, confirming both the scaling law in MS2 prediction and the practical value of π-MSNet. The improvement was particularly pronounced for TMT-labeled peptides carrying TMTpro 16plex modifications at both the N terminus and lysine (K) residues, with the PCC90 on unseen peptides increasing from 0 to 0.83, whereas the available TMT-specific Prosit model supports only N-terminal TMTpro 16plex modifications. Furthermore, using π- MSNet-retrained AlphaPeptDeep for PSM rescoring outperformed the original model, identifying on average 58 additional unique peptides per run at 1% FDR, with the median PCC increasing from 0.94 to 0.96.

**Fig. 3.**
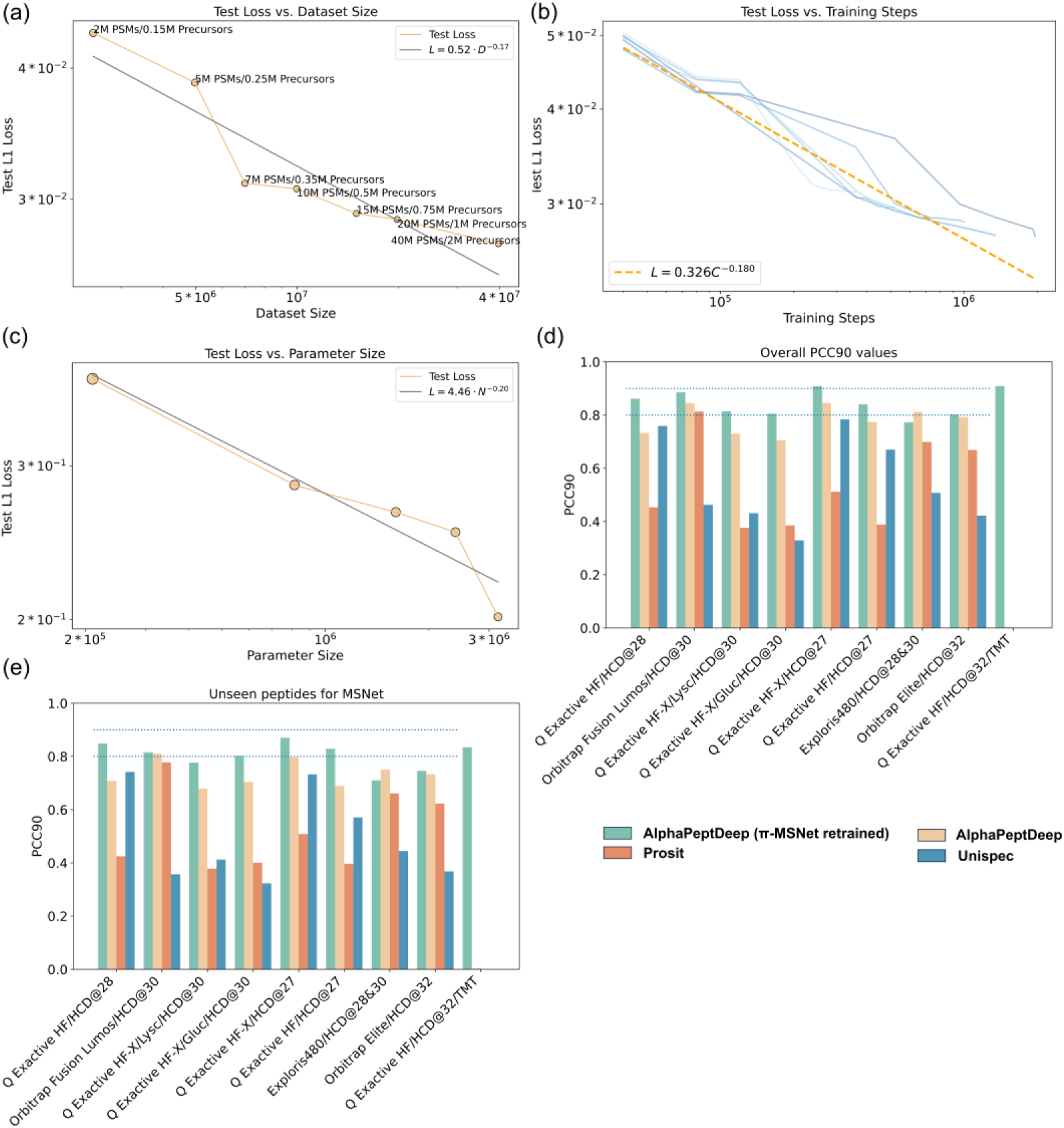
| **Scaling laws in MS2 intensity prediction and models benchmarking**. Modeling performance improves smoothly as we increase (a) the dataset size (b) the training steps and (c) the model size used for training AlphaPeptDeep. In (b), lines represent different training dataset sizes, with darker colors indicating larger datasets. Empirical performance follows a power-law relationship with each factor when not bottlenecked by the others. (d) Overall MS2 prediction accuracy (PCC90, the percentage of PCC values>0.9) of the three published models and π-MSNet-retrained AlphaPeptDeep on different test datasets. Dataset names are on the x-axis. (e) MS2 prediction accuracy for unseen peptides, defined as peptides not present in the training set across different models. Two dashed lines indicate PCC90 values of 0.8 and 0.9, respectively.

At the chromatographic level, RT prediction captures peptide elution behavior. In RT prediction, we retrained GPTime^25^, AutoRT^26^, and DeepLC using all available RT data in π-MSNet and evaluated their performance on independent test sets using R² (**Figure 4a–f**). We considered two data-splitting strategies: (i) grouping peptides by N-terminal amino acid, with peptides beginning with a specific amino acid assigned to the test set and all remaining peptides used for training; and (ii) grouping peptides by species, with peptides from a specific species assigned to the test set and peptides from all other species used for training. Once the training set contained approximately 10,000 peptides, further increases in its size resulted in only limited performance gains. We also examined the relationship between MAE and training set size. As shown in **Figure 4** **(a–f)**, R² curves exhibit a clear inflection point when the training set exceeds approximately 10,000 peptides. However, the MAE results in **Figure 4** **(g–j)** continue to decrease as the training set grows. Thus, the RT prediction task exhibits some degree of scaling behavior. Although simply increasing the number of training peptides yielded only marginal gains, the species-based evaluation suggests that the composition of the training data may also influence prediction performance. All three tools performed worst when *Homo sapiens* or *Mus musculus* peptides were used for testing, even though these two species accounted for substantially more peptides and projects than the other species. Together, these observations suggest that further improvements in RT prediction may require more informative strategies for grouping and selecting training data—for example, based on physicochemical properties or detailed chromatographic conditions—to reduce redundancy within groups and increase the diversity of the training data.

**Fig. 4.**
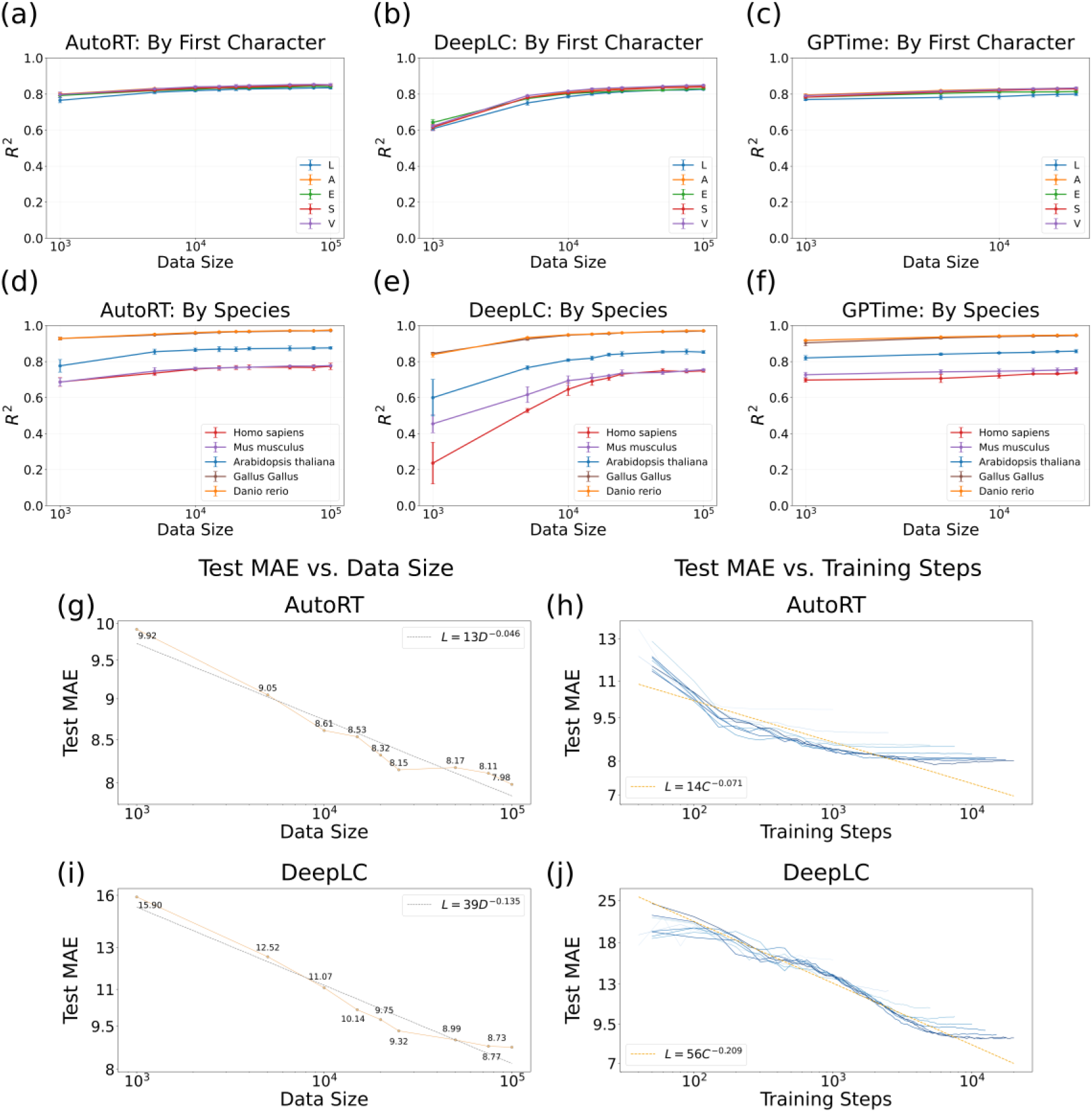
| **RT prediction performance of AutoRT, DeepLC, and GPTime using π-MSNet data.** (a–c) show the results of the leave-one-N-terminal-amino-acid- group-out evaluation for AutoRT, DeepLC, and GPTime, respectively, whereas (d–f) show the corresponding results of the leave-one-species-group-out evaluation. For each held-out group, peptides from that group were used exclusively for testing, while peptides from all remaining groups constituted the training set. In each of three independent rounds, 10,000 peptides were randomly sampled from the held-out group to form the test set. Each point represents the mean R² across the three rounds, and error bars indicate the standard deviation. R² denotes the coefficient of determination between the predicted and observed RTs in the held-out test set. (g,h), AutoRT scaling-law results for held-out Homo sapiens peptides; (i,j), corresponding DeepLC results. The left plot for each software shows converged MAE as a function of training- set size, whereas the right plot shows test MAE across training steps for different training-set sizes, with darker loss curves indicating larger training-set sizes. Dashed lines indicate log-log power-law fits. All scaling-law models were trained from scratch using nested subsets from the non-Homo sapiens training pool and evaluated on a randomly sampled 1,000-peptide Homo sapiens test set.

De novo peptide sequencing, which identifies peptides from tandem mass spectrometry data without relying on reference protein databases, holds significant value in complex biological scenarios where constructing accurate reference databases is challenging, such as immunopeptidomics, antibody sequencing, and metaproteomics. In recent years, the explosion of deep learning technologies has brought revolutionary breakthroughs to de novo sequencing. Notably, π-HelixNovo27 is a deep-learning-based de novo peptide sequencing model previously developed by our group, which achieved state-of-the-art performance on a nine-species benchmark dataset. Here, to demonstrate the utility of π-MSNet and evaluate the benefits of high-quality training data, we retrained π-HelixNovo using a subset of π-MSNet and refer to this retrained model as π-HelixNovo-MSNet.

We compared π-HelixNovo-MSNet with the optimal model provided by π- HelixNovo (denoted π-HelixNovo-raw) on the widely used “Nine-species” dataset^28^, using the leave-one-specie-out strategy, in which the rice bean dataset served as the test set and was held out from training. We evaluated two models on same test sets, as shown in **Figure 5a**. The results demonstrate that π-HelixNovo-MSNet outperforms π-HelixNovo-raw in peptide recall across all tested species, achieving a 39.1% relative increase in average recall per dataset.

**Fig. 5.**
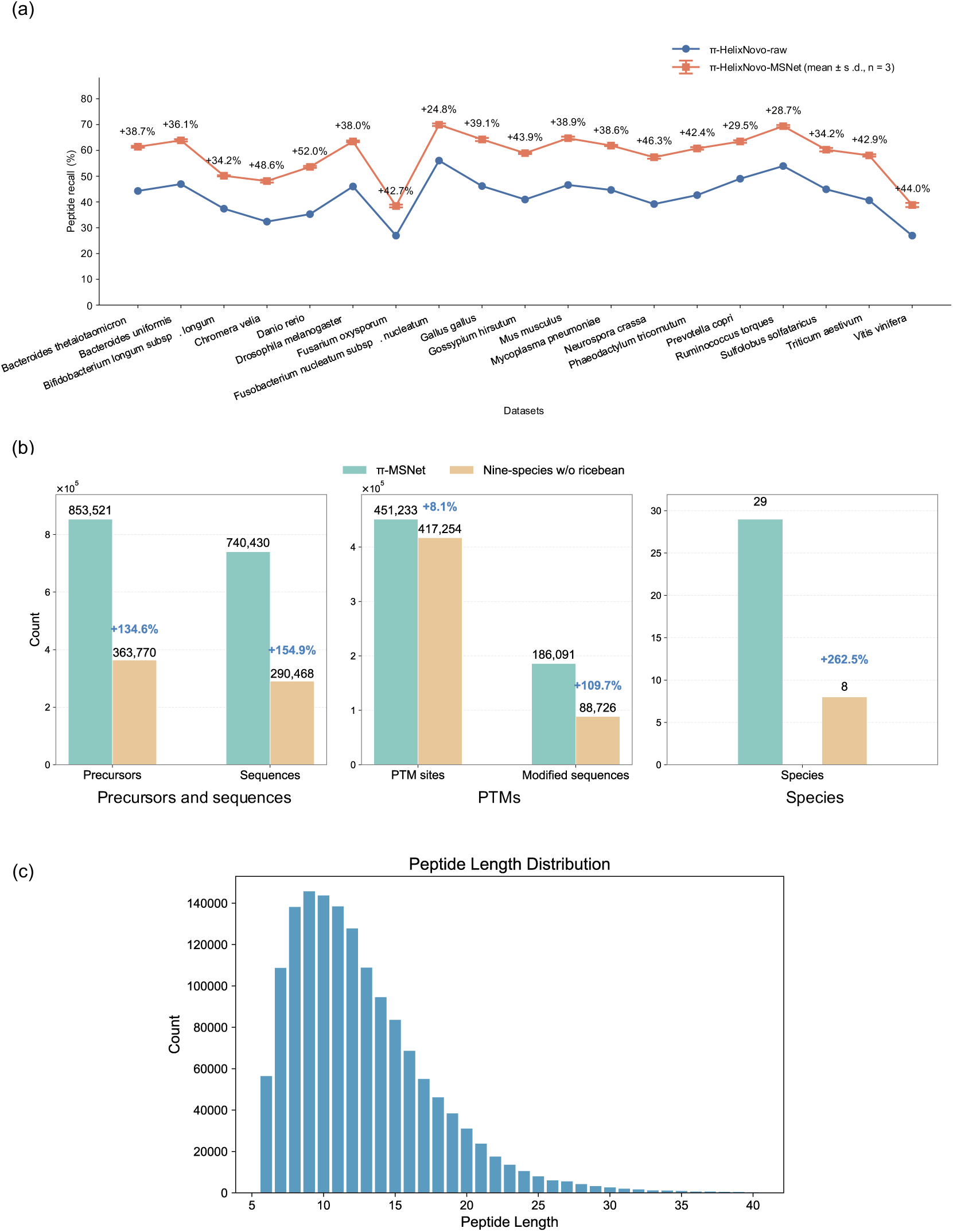
| **Performance enhancement of de novo peptide sequencing using π-MSNet with the same training data size (1.49 million).** (a) Peptide recall comparison between π-HelixNovo-MSNet and π-HelixNovo-raw across multi- species datasets. (b) Comparison of dataset characteristics between π-MSNet training set (1.49 million) and the “Nine-species” dataset (excluding the rice bean dataset, 1.49 million). (c) Peptide length distribution in the down-sampled 1.49 million training set from π-MSNet.

Given that the training scales, training parameters, configurations, and test sets were identical for both models, we attribute this performance enhancement primarily to the intrinsic qualities of the π-MSNet dataset. We further compared the characteristics of π-MSNet and the “Nine-species” dataset (excluding the rice bean data) (Figure 5b). First, it is worth noting that, for a fair comparison, we subsampled the π-MSNet dataset to keep the same sample scale (1.49 million PSMs) with π-HelixNovo-raw training set. In this circumstance π-MSNet exhibits substantial increases in precursor and peptide diversity of 134.6% and 154.9%, respectively, compared to the “Nine-species” dataset. The core objective of de novo sequencing is to learn generalized peptide fragmentation patterns; thus, a broader repertoire of peptides and precursors in the training data enables the model to capture more comprehensive fragmentation rules, significantly bolstering its generalization capability. Furthermore, PTMs typically exist at low abundances and pose a major challenge for de novo sequencing. π-MSNet contains 8.1% more modification sites and 109.7% more modified peptides than the “Nine-species” dataset (considering methionine oxidation, cysteine carbamidomethylation, and acetylation). This higher proportion of modified data enhances the model’s sensitivity in identifying low-abundance PTMs. Finally, the inclusion of a wider taxonomic range (29 vs 8) in π-MSNet intrinsically strengthens the model’s cross-species sequencing capacity, which is vital for applications in complex fields like metaproteomics.

Additionally, we analyzed the peptide length distribution within the π-MSNet dataset (**Figure 5c**). The lengths range from 6 to 40 amino acids, with an average of 12.6, spanning 35 distinct length categories. Notably, π-MSNet implements rigorous quality control regarding peptide length: the majority of peptides are concentrated between 7 and 15 amino acids, minimizing the presence of excessively short or long sequences (in contrast, the “Nine-species” dataset spans lengths from 5 to 65). This stringent length filtering (peptide length from 6 to 40) ensures that the training data falls strictly within the optimal detection range of mass spectrometry, mitigating the interference of outliers during model training. This high structural diversity facilitates robust model generalization across spectra of varying peptide lengths. In summary, characterized by its high quality and extensive diversity, the π-MSNet dataset holds a great promise for driving further performance breakthroughs in de novo sequencing models. Furthermore, by leveraging larger data subsets from the π-MSNet repository, the research community can expect to unlock even more substantial improvements in model accuracy and generalization. Detailed procedures for dataset curation, model training, and downstream analyses are provided in the **Methods** section.

Besides above three tasks, we also explored collisional cross section (CCS) prediction with our data resource. Retraining the AlphaPeptDeep CCS model on progressively larger subsets of PXD019086 revealed a power-law decrease in test loss with increasing training data, with larger datasets benefiting from additional training steps. In contrast, gains from increasing model size diminished beyond approximately 3 million parameters.

To enable the user-friendly utilization of retrained models, we developed π- MSNet agent for mass spectrometry data analysis. Users can interact conversationally for MS2 intensity prediction, DIA library construction, retention time prediction, de novo sequencing and visualization. We evaluated the agent in a 20-case benchmark covering these four proteomics tasks. Three browser- use models, three non-expert users and three expert users each completed the 20 cases, yielding 180 runs. Across 180 benchmark runs, which were successfully completed, 97.8% of the final results were consistent with expert-generated references, and agent-assisted workflows were faster than manual execution in 150 runs (83.3%), excluding one-time deployment, while 175 runs (97.2%) required only two or three interaction turns and eight (4.4%) required a retry (Figure 6). These results demonstrate that π-MSNet provides an efficient and reliable conversational interface for model invocation, proteomics analysis and result interpretation. The agent is publicly available at https://msnet.ncpsb.org.cn/ or https://msagent.wisdomeyes.cn.

**Fig. 6.**
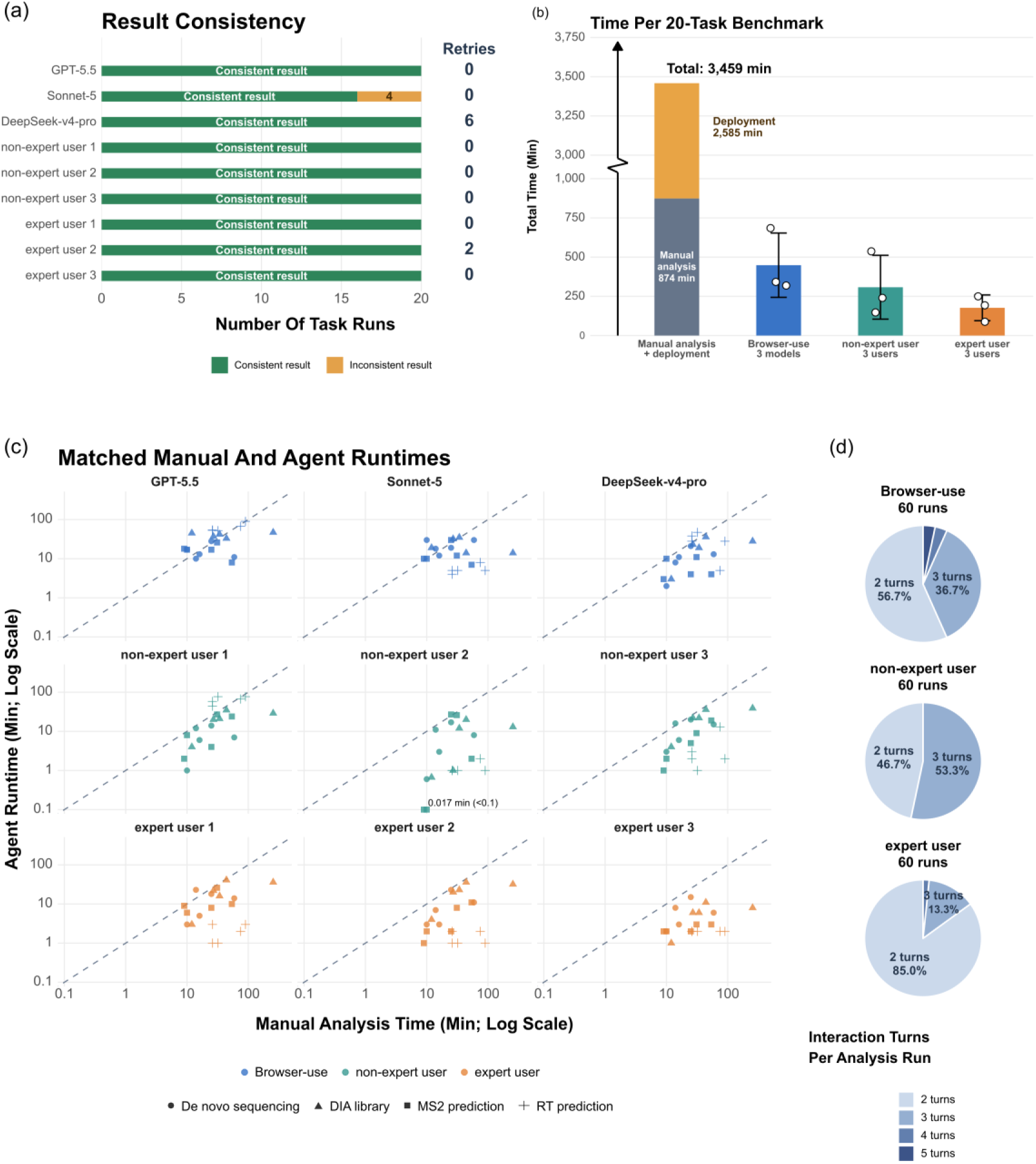
| **Benchmarking agent performance efficiency and reliability.** (a) Result consistency and number of retries across the 20 benchmark cases for each model or user. Consistency was assessed against the results generated by the expert reference workflow. Stacked bars indicate consistent and inconsistent results, and the adjacent column indicates the number of runs requiring a retry before the final result was accepted. (b) Total time required to complete the 20-case benchmark. Bars show the mean summed runtime ± sample SD across three browser-use models, three non-expert users, and three expert users; open circles indicate individual totals. Manual analysis time is separated into script development/analysis and one-time deployment. (c) Matched manual and agent-assisted runtimes for all 180 analysis runs. Colors denote interaction groups and shapes denote workflows. Both axes are logarithmic. The dashed identity line indicates equal runtime; points below the line indicate faster agent-assisted execution. (d) Distribution of the number of interaction turns with the π-MSNet agent for browser-use agents (n = 60), non- expert users (n = 60), and expert users (n = 60).

Overall, π-MSNet aims at resolving the long-standing challenge of unstructured, poorly annotated and costly MS data by delivering the a large-scale and rigorously curated proteomics portal. Leveraging our high-quality resource, we empirically reproduced scaling laws for specific tasks, revealing that enhancing peptide diversity is the key factor for improving the predictive performance of MS-based models. π-MSNet not only enables rigorous performance evaluation and optimization of MS models but also serves as an essential platform for downstream task development, ultimately advancing data-driven discovery in proteomics.

## Discussion

The accelerating adoption of deep learning in proteomics has introduced a critical new requirement for data infrastructure: large-scale, AI-ready, and continuously reprocessed data resource for robust model training and evaluation. Although neural network architectures for MS2 prediction, RT prediction, and de novo peptide sequencing have advanced rapidly, their performance and generalizability remain fundamentally constrained by the scale, diversity, and consistency of available training data. By systematically reanalyzing and harmonizing tens of thousands of public LC-MS/MS runs within a unified workflow and a machine learning-tailored format, π-MSNet establishes a scalable, continuously reprocessed data resource for reproducible model training and evaluation. Rather than introducing a new predictive algorithm, π-MSNet addresses an upstream bottleneck: the lack of standardized, AI-ready, continuously updated datasets capable of supporting large-scale computational development. Building on this data resource and retrained models, the π-MSNet agent extends π-MSNet to an interactive framework for proteomics data analysis, supporting tasks including MS2 prediction, RT prediction, DIA library prediction and de novo peptide sequencing. Recent agentic systems such as AstroAgents29 and MSAgent30 have explored mass spectrometry data analysis in metabolomics and other contexts, whereas the π-MSNet Agent is specifically designed for processing large-scale proteomics data and corresponding computational tasks using the models retrained on π-MSNet data.

The architecture of π-MSNet is designed to address the rapid growth and diversification of public MS data. Static datasets inevitably become outdated as acquisition technologies, instrument configurations and biological applications change. By providing incremental updates, π-MSNet provides a continuously reprocessing infrastructure rather than a fixed benchmark snapshot. Sustained community participation, transparent curation practices and consistent metadata schemas will be critical to ensure long-term utility and interoperability.

Our retraining experiments suggest that increasing data diversity and scale improves cross-dataset robustness and reduces sensitivity to experiment-specific biases. These observations are consistent with broader trends in machine learning, where performance scaling often depends on both dataset size and heterogeneity. The availability of large-scale, multi-instrument MS data creates opportunities to investigate scaling behaviors, transfer learning strategies and foundation model paradigms tailored to proteomics. Systematic exploration of how model capacity and training data scale interact may provide quantitative guidance for future architecture design.

Despite these advances, π-MSNet still has limitations. First, it does not encompass the full spectrum of proteomics data types yet, such as cross-linking mass spectrometry (XL-MS). Ongoing incremental updates are expected to further expand both the scale and diversity of the resource. Second, PTM coverage remains limited due to constraints in both experimental techniques (e.g., PTM enrichment) and computational identification methods, despite the inclusion of 23 modification types in the current release. Future development of π-MSNet will focus on expanding PTM diversity through the integration of additional datasets and the adoption of advanced identification strategies, such as open modification search approaches.

## Methods

### Data Collection and Preparation

The initial construction of the π-MSNet repository used over 40 TB of MS data from 127 public datasets, including PRIDE Archive, iProX and the in-house data from the π-HuB project. All raw data is available at the corresponding repositories. The reanalyzed files are also available at https://msnet.ncpsb.org.cn and https://portal.quantms.org. All proteomics datasets are annotated in SDRF-Proteomics sample metadata format for further analysis.

## Processing workflow for π-MSNet datasets

The processing workflow of π-MSNet is based on quantms v1.3 and quantmsdiann v2.1.0, both of which are cloud-based proteomics identification and quantification workflow. MSGF+ and Comet were selected as search engines in quantms for DDA data. Percolator calculated a posterior error probability (PEP) for each PSM. Then, the ConsensusID tool combined the PSMs from multiple search engines into a final score for each PSM. After ConsensusID, file-wide PSM-level q values were taken from Percolator. For quantmsdiann analyses, the workflow was configured to use DIA-NN v2.5.1.

The tandem mass spectra from different species were searched against their respective reference proteome databases. The search parameters were obtained from the original papers and stored in SDRF metadata files, including modification, precursor mass tolerance and fragment mass tolerance.

For all the datasets in π-MSNet, a stringent FDR at 1% was applied for each project at the PSM level. Specifically, for phosphorylation datasets, a false localization rate of 0.01 was further applied after PSM filtering. All matched peaks (b+, b++, y+, y++, with and without neutral losses) were annotated for each PSM from DDA data using the spectrum-utils package, with different fragment mass tolerance settings applied during annotation. The final identification results were stored in QPX format.

## Training and benchmarking MS2 intensity model

For MS2 intensity model training and benchmarking, all train and test datasets are listed. We included spectra obtained using nine instruments, four enzyme cleavages and eight normalized collision energy settings (NCEs). The matched b and y ions were annotated using spectrum-utils package for each PSMs after quantms identification. Only b+, b++, y+ and y++ ions were considered for MS2 intensity prediction task, as common methods focus on training their networks to accurately predict intensities of b and y fragments. For consistency, we used the same peak-matching tolerances used in quantms. If no matching experimental peak was found, then the corresponding experimental intensity was set to zero. The matched b and y fragment intensities in the training and testing datasets were normalized by dividing by the highest matched intensity for each spectrum. For scaling-law validation, the π-MSNet training parameters were: epoch=100, warmup epoch=20, learning rate (lr)=1e-5, dropout=0.1. For model retraining, we followed a similar multi-stage training strategy. In the first stage, AlphaPeptDeep was trained on DDA LFQ data with the following parameters: epoch=100, warmup epoch=20, lr=1e−5, and dropout=0.1. In the second stage, the model was further trained on TMT-labeled data with epoch=20, warmup epoch=5, lr=1e−5, and dropout=0.1. The peptides from the training datasets were not included in tested datasets for scaling law experiments. L1 loss was used for all training phases. Except for retraining AlphaPeptDeep on π-MSNet, each model was accessed via Koina API for prediction.

## Training and benchmarking retention time model

Peptide RT values of all π-MSNet projects were normalized to a 0-100 min range within each raw MS run. The data were subsequently deduplicated at the peptide level, and the mean normalized RT was used as the final prediction target. The resulting peptide-level data were used to train and evaluate AutoRT, DeepLC, and GPTime. AutoRT (v2.0.0-beta) and DeepLC (v3.1.13) were trained from scratch using their default 10-model and 3-model ensemble configurations, respectively. GPTime was run using its default repository workflow with GPTIME_OPTIMIZE_RESTARTS=1.

To prevent peptide-level data leakage, leave-one-group-out splits were constructed independently by peptide first character and species, and were applied consistently to all three software packages. The five first-character groups with the largest peptide counts (L, A, E, S, and V) and the five species with the largest peptide counts (Homo sapiens, Mus musculus, Arabidopsis thaliana, Gallus gallus, and Danio rerio) were selected as held-out test groups respectively. In each of three independent rounds, 10,000 peptides were randomly sampled from the held-out group as the test set, whereas the training data contained peptides only from the remaining groups. The three test sets generated for each held-out group were non-overlapping. Nested training subsets of 1,000, 5,000, 10,000, 15,000, 20,000, 25,000, 50,000, 75,000, and 100,000 peptides were evaluated. Because the computational and memory requirements of GPTime increased rapidly with training-set size, GPTime was evaluated only up to 25,000 training peptides. Predictive performance was quantified using the coefficient of determination (R2), reported as the mean and standard deviation over the three rounds.

Scaling-law experiments were performed with AutoRT and DeepLC across the five species with the largest total peptide counts. For each species, 1,000 peptides were randomly sampled as the test set, and nested training subsets of 1,000, 5,000, 10,000, 15,000, 20,000, 25,000, 50,000, 75,000, and 100,000 peptides were constructed from the remaining species. Test-set MAE was evaluated throughout model training. Each test-loss trajectory was smoothed using a progressively expanding moving-average window, and the converged MAE at each training-set size was selected for scaling-law analysis. Power-law relationships between MAE and either training steps or training-set size were fitted in log-log space.

## Retraining and evaluation of π-HelixNovo

The files in PXD014877 dataset were randomly partitioned in a 7:3 ratio to construct the training and test sets. 1.49 million PSMs (same size as the original π-HelixNovo model’s training set) were randomly sampled three times from the training set to retrain π-HelixNovo-MSNet. To establish a baseline for performance comparison, referred to as π-HelixNovo-raw in this study, we used the original pre-trained model weights provided by the authors of π-HelixNovo and evaluated this model on the same test set as π-HelixNovo-MSNet. The source code for the de novo sequencing model was obtained directly from the official π-HelixNovo GitHub repository (https://github.com/PHOENIXcenter/pi-HelixNovo).

To rigorously prevent data leakage and ensure an unbiased evaluation, we enforced strict peptide-level deduplication: any spectra in the test set whose corresponding peptide sequences appeared in the training set were completely discarded. For hyperparameter tuning and model selection, the rice bean dataset—derived from the widely adopted “Nine-species” benchmark dataset (MassIVE: MSV000081382)—was employed as the validation set. Similarly, to maintain the absolute independence of the test data, we excluded any spectra from this validation set that shared peptide sequences with the test set.

During the retraining process of π-HelixNovo-MSNet, all hyperparameter configurations were kept strictly identical to those described in the original π- HelixNovo publication to ensure a fair and controlled comparison. The model checkpoint that achieved the lowest validation loss on the rice bean dataset was selected for final benchmarking on the multi-species test sets. Performance evaluation was conducted using the identical metrics adopted by π-HelixNovo. All benchmarking inferences were executed under the model’s standard evaluation mode to ensure deterministic outputs.

## Benchmarking the π-MSNet agent

We evaluated the agent using 20 representative cases: five each for MS2 intensity prediction, DIA library prediction, de novo sequencing and retention time prediction. Three browser-use models (GPT-5.5, Sonnet-5 and DeepSeek- v4-pro), three non-expert users and three expert users each completed all 20 cases, for 180 runs in total. The browser-use models interacted with the agent through its web interface as simulated users. We compared final outputs with expert-generated reference results and recorded analysis time, interaction turns and whether a retry was needed before accepting the final result. The time required for expert manual analysis was used as the reference; matched runtime comparisons excluded one-time environment setup and model deployment.

## Data availability

All datasets in this portal are available at https://msnet.ncpsb.org.cn and https://portal.quantms.org.

## Code availability

Open access source codes for loading π-MSNet datasets into PyTorch and TensorFlow, as well as retrained models, are available at https://github.com/PHOENIXcenter/pi-MSnet^31^.

## Acknowledgements

This work is supported by the National Key Research and Development Program of China (2025YFA1309300), the National Natural Science Foundation of China (32688201), the CAMS Innovation Fund for Medical Sciences (CIFMS, 2019-I2M-5-063) and the research program of the State Key Laboratory of Medical Proteomics (SKLP-Y202401). We thank Jingwen Ye, Huanjia Liang, Zhitao Gu, and Juan Chen for their contributions to improving the π-MSNet data portal, as well as Bolun Wang and Zicheng Ma for their assistance in compiling the retention time data and evaluating the RT prediction tools.

## Author contributions

The study was designed by C.C. Supervision was provided by F.H., Y.P.-R., L.X. and C.C. Data curation, reanalysis and methodology development were carried out by C.D., Y.L., T.L, M.W. and S.R. π-MSNet web portal was developed by Y.Q. and H.X. π-MSNet agent was developed and updated by Q.Z and Y.C. and X.H. The original draft was written by C.D., Y.L., and T.L. Writing review and editing were done by Y.Z., M.B., T.S., F.H., Y.P.-R., L.X. and C.C. All authors approved the final manuscript.

## Competing interests

The authors declare no competing interests.

## Notes

https://msnet.ncpsb.org.cn

https://portal.quantms.org

### Competing Interest Statement

The authors have declared no competing interest.

### Summary of Updates

1.MSNet has been expanded substantially in both scale and experimental diversity, with the number of PSMs increasing from approximately 501 million at initial submission to over 737 million in this revision. We further incorporated DIA, TMT labeled, SILAC labeled and metaproteomics datasets. The resource now comprises 11 million precursors derived from 45,583 LC MS/MS runs spanning 12 instrument types and 55 species, including 152 million DIA PSMs and 76.6 million DDA TMT-labeled PSMs. 2.We have added operational evidence for the continuously reprocessing architecture of MSNet, including benchmarks of large-scale reanalysis, incremental data releases during the review and revision period and a standardized data-upload interface for community-contributed datasets. 3.We have strengthened the evaluation of AI models through more stringent peptide- and species-level data-splitting strategies and additional scaling analyses for MS2 intensity, retention time and collisional cross section prediction. 4.For de novo peptide sequencing, we have performed size-matched comparisons and a controlled precursor-diversity experiment to distinguish the contribution of data diversity from that of training-set size. 5.We have substantially expanded the evaluation of the MSNet agent through a 180-run benchmark involving browser-use models (imitating human users with large language models), non-expert users and expert users, assessing result consistency to the reference, analysis efficiency, interaction complexity and retry behavior.

## References

1. Mann, M., Kumar, C., Zeng, W.-F. & Strauss, M. T. Artificial intelligence for proteomics and biomarker discovery. Cell Systems 12, 759–770 (2021).

2. Zeng, W.-F. et al. AlphaPeptDeep: a modular deep learning framework to predict peptide properties for proteomics. Nat Commun 13, 7238 (2022).

3. Bouwmeester, R., Gabriels, R., Hulstaert, N., Martens, L. & Degroeve, S. DeepLC can predict retention times for peptides that carry as-yet unseen modifications. Nat Methods 18, 1363–1369 (2021).

4. Lautenbacher, L. et al. Koina: Democratizing machine learning for proteomics research. Nat Commun 16, 9933 (2025).

5. Yilmaz, M. et al. Sequence-to-sequence translation from mass spectra to peptides with a transformer model. Nat Commun 15, 6427 (2024).

6. Zhang, X. et al. π-PrimeNovo: an accurate and efficient non-autoregressive deep learning model for de novo peptide sequencing. Nat Commun 16, 267 (2025).

7. Deutsch, E. W. et al. The ProteomeXchange consortium in 2026: making proteomics data FAIR. Nucleic Acids Research 54, D459–D469 (2026).

8. Perez-Riverol, Y. et al. The PRIDE database at 20 years: 2025 update. Nucleic Acids Res 53, D543–D553 (2025).

9. Ma, J. et al. iProX: an integrated proteome resource. Nucleic Acids Research 47, D1211–D1217 (2019).

10. Wilkinson, M. D. et al. The FAIR Guiding Principles for scientific data management and stewardship. Sci Data 3, 160018 (2016).

11. Kaplan, J., et al. Scaling Laws for Neural Language Models. Preprint at 10.48550/arXiv.2001.08361 (2020).

12. Dens, C., Adams, C., Laukens, K. & Bittremieux, W. Machine Learning Strategies to Tackle Data Challenges in Mass Spectrometry-Based Proteomics. J. Am. Soc. Mass Spectrom. jasms.4c00180 (2024).

13. Wen, B. & Noble, W. S. A multi-species benchmark for training and validating mass spectrometry proteomics machine learning models. Sci Data 11, 1207 (2024).

14. Wang, M. et al. Assembling the Community-Scale Discoverable Human Proteome. Cell Systems 7, 412–421.e5 (2018).

15. He, F. et al. π-HuB: the proteomic navigator of the human body. Nature 636, 322–331 (2024).

16. Dai, C. et al. A proteomics sample metadata representation for multiomics integration and big data analysis. Nat Commun 12, 5854 (2021).

17. Dai, C. et al. quantms: a cloud-based pipeline for quantitative proteomics enables the reanalysis of public proteomics data. Nat Methods 21, 1603–1607 (2024).

18. Yue, Q. et al. quantmsdiann: a scalable SDRF-driven DIA-NN workflow for reanalysis of single-cell, spatial, and bulk proteomics datasets. Research Square (2026).

19. Kim, S. & Pevzner, P. A. MS-GF+ makes progress towards a universal database search tool for proteomics. Nat Commun 5, 5277 (2014).

20. McGann, C. D. et al. Comet Fragment-Ion Indexing for Enhanced Peptide Sequencing. J. Proteome Res. 24, 3715–3721 (2025).

21. Käll, L., Canterbury, J. D., Weston, J., Noble, W. S. & MacCoss, M. J. Semi-supervised learning for peptide identification from shotgun proteomics datasets. Nat Methods 4, 923–925 (2007).

22. Demichev, V., Messner, C. B., Vernardis, S. I., Lilley, K. S. & Ralser, M. DIA-NN: neural networks and interference correction enable deep proteome coverage in high throughput. Nat Methods 17, 41–44 (2020).

23. Gessulat, S. et al. Prosit: proteome-wide prediction of peptide tandem mass spectra by deep learning. Nat Methods 16, 509–518 (2019).

24. Lapin, J., Yan, X. & Dong, Q. UniSpec: Deep Learning for Predicting the Full Range of Peptide Fragment Ion Series to Enhance the Proteomics Data Analysis Workflow. Anal. Chem. 3c02321 (2024).

25. Maboudi Afkham, H., Qiu, X., The, M. & Käll, L. Uncertainty estimation of predictions of peptides’ chromatographic retention times in shotgun proteomics. Bioinformatics 33, 508–513 (2017).

26. Wen, B., Li, K., Zhang, Y. & Zhang, B. Cancer neoantigen prioritization through sensitive and reliable proteogenomics analysis. Nat Commun 11, 1759 (2020).

27. Yang, T. et al. Introducing π-HelixNovo for practical large-scale de novo peptide sequencing. Briefings in Bioinformatics 25, bbae021 (2024).

28. Tran, N. H., Zhang, X., Xin, L., Shan, B. & Li, M. De novo peptide sequencing by deep learning. Proc. Natl. Acad. Sci. U.S.A. 114, 8247–8252 (2017).

29. Saeedi, D., Buckner, D., Aponte, J. C. & Aghazadeh, A. AstroAgents: A Multi-Agent AI for Hypothesis Generation from Mass Spectrometry Data. Preprint at 10.48550/ARXIV.2503.23170 (2025).

30. Li, Y. et al. MSAgent: An Evidence Grounded Agentic Framework for LLM-driven Scientific Exploration in Mass Spectrometry-based Metabolomics. Preprint at 10.64898/2026.04.22.720103 (2026).

31. Dai, C. et al. π-MSNet: A large-scale, AI-ready and continuously reprocessed proteomics data resource v0.0.4. Zenodo 10.5281/zenodo.22975527 (2026).

