## Supplementary Information for "π-MSNet: A large-scale, AI-ready and continuously reprocessed proteomics data resource"

Supplementary Note 1: The π-MSNet continuously reprocessed datasets.

A total of 127 datasets were reanalyzed using quantms v1.3 and quantmsdiann v2.1.0 against UniProt protein sequence database, only the SwissProt (reviewed proteins) without isoforms information^1,2^. For quantmsdiann analyses, the workflow was configured to use DIA-NN v2.5.1^3^. All qpx results have been deposited in <https://msnet.ncpsb.org.cn> and https://portal.quantms.org/collections. Supplementary Table 1 shows the list of datasets reanalyzed including species, instrument, enzyme, ms run, and peptide spectrum matches for each dataset until 09.10.2026.

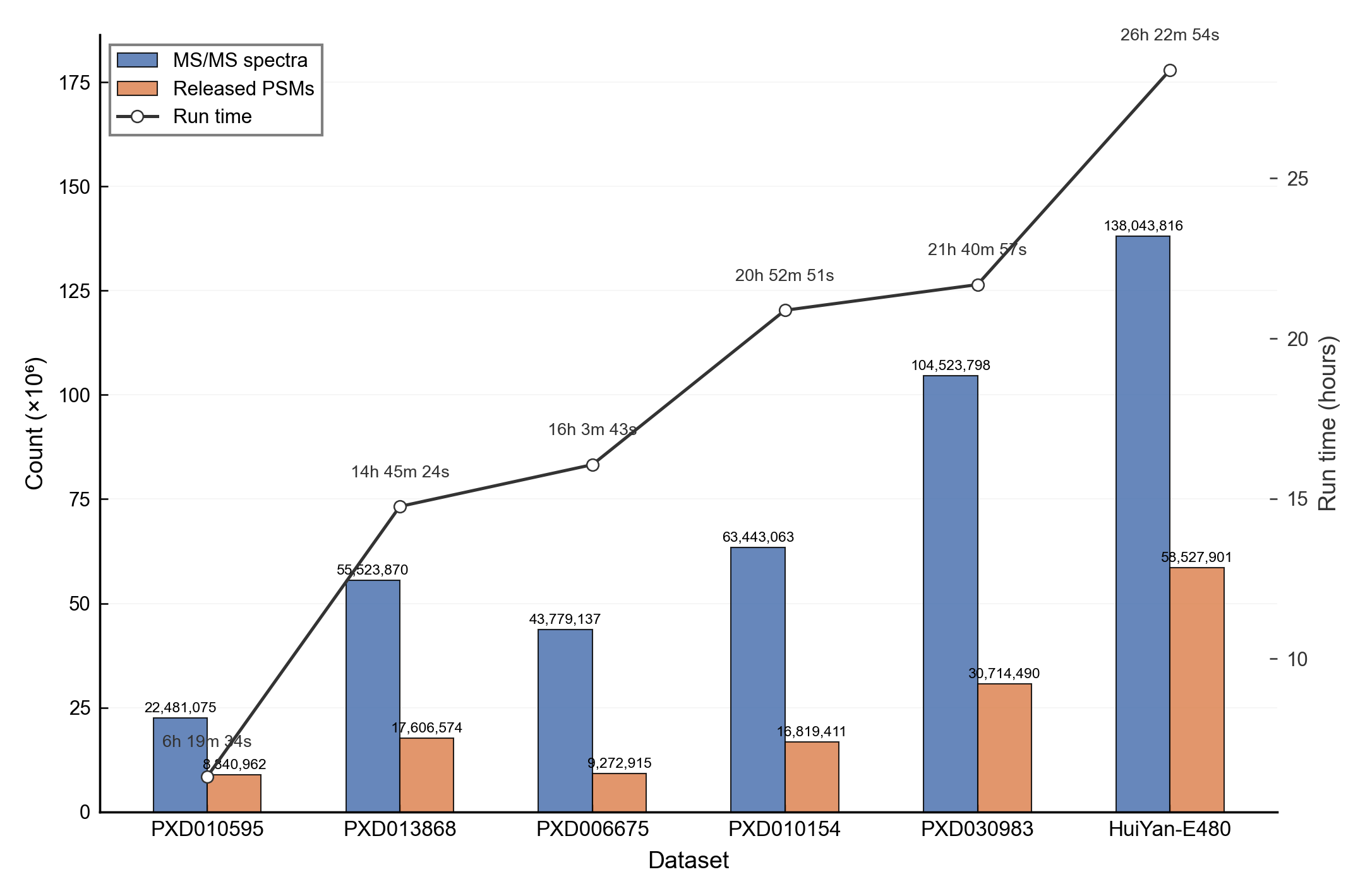

Supplementary Figure 1: Actual quantms pipeline run time across datasets of varying data volumes in an HPC cluster with 500 CPU cores. The plotted lines together with blue bars show the relationship between run time and total MS2 spectra acquired, while the orange bars indicate the number of released PSMs. Each point on the lines represents a benchmark dataset, labeled by project accession. These plots illustrate the computational time required in practice to process MS2 data of different scales and the corresponding PSM yield after FDR filtering. PSM: peptide-spectrum match.

Supplementary Table 1: Datasets reanalyzed using quantms and quantmsdiann.

| *Accession* | *Species* | *Instrument* | *Enzyme* | #*MS run* | #PSM |
| --- | --- | --- | --- | --- | --- |
| HuiYan-E480 | *Homo sapiens* | Orbitrap Exploris 480 | Trypsin | 40 | 58527091 |
| HuiYan-Bat | *Rhinolophus sinicus* | timsTOF | Trypsin | 763 | 2334922 |
| IPX0000937001 | *Homo sapiens* | Orbitrap Exploris 480 | Trypsin | 1344 | 24924726 |
| IPX0000937003 | *Homo sapiens* | Orbitrap Fusion | Trypsin | 585 | 6551233 |
| IPX0001289001 | *Homo sapiens* | Orbitrap Fusion | Trypsin | 1139 | 17387674 |
| IPX0001590001 | *Homo sapiens* | Orbitrap Fusion Lumos | Trypsin | 659 | 7488098 |
| IPX0001804001 | *Homo sapiens* | Q Exactive Plus | Trypsin | 2136 | 23982246 |
| IPX0001804002 | *Homo sapiens* | Orbitrap Fusion | Trypsin | 451 | 1426890 |
| IPX0002031000 | *Homo sapiens* | Orbitrap Fusion | Trypsin | 724 | 27013882 |
| IPX0004073001 | *Homo sapiens* | Q Exactive HF-X | Trypsin | 120 | 1018214 |
| PXD000561 | *Homo sapiens* | Orbitrap Fusion Lumos | Trypsin | 2187 | 10334261 |
| PXD000612 | *Homo sapiens* | Q Exactive HF | Trypsin | 231 | 6416764 |
| PXD000865 | *Homo sapiens* | Q Exactive | Chymotrypsin | 168 | 148896 |
| PXD000865 | *Homo sapiens* | LTQ Orbitrap Velos | Lys-C | 120 | 529187 |
| PXD000865 | *Homo sapiens* | LTQ Orbitrap Elite | Trypsin | 767 | 4990620 |
| PXD002137 | *Homo sapiens* | LTQ Orbitrap Velos | Trypsin | 192 | 18775812 |
| PXD002395 | *Homo sapiens* | Q Exactive | Trypsin | 198 | 3796473 |
| pxd002767 | *Solanum tuberosum* | LTQ Orbitrap Velos | Trypsin | 297 | 2666095 |
| PXD002854 | *Homo sapiens* | Q Exactive | Trypsin | 174 | 548653 |
| PXD003903 | *Homo sapiens* | Q Exactive HF | Trypsin | 335 | 7325444 |
| PXD003947 | *Homo sapiens* | Q Exactive Plus | Trypsin | 103 | 740197 |
| PXD004242 | *Homo sapiens* | Q Exactive HF | Trypsin | 1289 | 5197467 |
| PXD004452 | *Homo sapiens* | Q Exactive HF | Gluc | 39 | 154347 |
| PXD004452 | *Homo sapiens* | Q Exactive HF | Trypsin | 789 | 5495744 |
| PXD004732 | *Homo sapiens* | Orbitrap Fusion Lumos | Trypsin | 538 | 14298028 |
| PXD006225 | *Drosophila melanogaster* | Q Exactive Plus | Trypsin | 31 | 536276 |
| PXD006675 | *Homo sapiens* | Q Exactive HF | Trypsin | 585 | 9272915 |
| PXD008333 | *Homo sapiens* | Q Exactive Plus | Trypsin | 201 | 2036467 |
| PXD008722 | *Homo sapiens* | Q Exactive HF | Trypsin | 252 | 3642440 |
| PXD008840 | *Homo sapiens* | Orbitrap Fusion | Trypsin | 504 | 9733821 |
| PXD009348 | *Homo sapiens* | Q Exactive HF | Trypsin | 163 | 386933 |
| PXD009449 | *Homo sapiens* | Orbitrap Fusion Lumos | Trypsin | 26 | 201402 |
| PXD009449 | *Homo sapiens* | Orbitrap Fusion Lumos | Trypsin | 39 | 697583 |
| PXD009737 | *Homo sapiens* | Q Exactive HF-X | Gluc | 36 | 349187 |
| PXD009737 | *Homo sapiens* | Q Exactive HF-X | Lys-C | 36 | 430334 |
| PXD009737 | *Homo sapiens* | Q Exactive HF-X | Trypsin | 36 | 573528 |
| PXD010154 | *Homo sapiens* | Q Exactive Plus | Trypsin | 1364 | 16819411 |
| PXD010595 | *Homo sapiens* | Orbitrap Fusion Lumos | Trypsin | 384 | 8840962 |
| PXD010899 | *Homo sapiens* | Q Exactive | Trypsin | 1803 | 1570723 |
| PXD010957 | *Mus musculus* | Q Exactive | Trypsin | 1437 | 22686367 |
| PXD012131 | *Homo sapiens* | Orbitrap Fusion Lumos | Trypsin | 311 | 4563808 |
| PXD012174 | *Homo sapiens* | LTQ Orbitrap Velos | Trypsin | 1194 | 15621996 |
| PXD012636 | *Danio rerio* | Q Exactive HF | Trypsin | 58 | 98759 |
| PXD012636 | *Equus caballus* | Q Exactive HF | Trypsin | 124 | 134634 |
| PXD012636 | *Mus musculus* | Q Exactive HF | Trypsin | 120 | 1568361 |
| PXD012636 | *Sus scrofa domesticus* | Q Exactive HF | Trypsin | 120 | 417991 |
| PXD012636 | *Rattus norvegicus* | Q Exactive HF | Trypsin | 120 | 937801 |
| PXD012636 | *Homo sapiens* | Q Exactive HF | Trypsin | 89 | 1132652 |
| PXD013231 | *Homo sapiens* | Orbitrap Fusion Lumos | Trypsin | 105 | 468370 |
| PXD013868 | *Arabidopsis* | Q Exactive HF | Trypsin | 1435 | 17606574 |
| PXD014877 | *Bacteroides Fragilis* | Q Exactive HF-X | Trypsin | 1 | 51628 |
| PXD014877 | *Bacteroides Uniformis* | Q Exactive HF-X | Trypsin | 1 | 54455 |
| PXD014877 | *Bifidobacterium Longum Subsp Longum* | Q Exactive HF-X | Trypsin | 1 | 21831 |
| PXD014877 | *Chromera velia* | Q Exactive HF-X | Trypsin | 5 | 68644 |
| PXD014877 | *Clostridium Bolteae* | Q Exactive HF-X | Trypsin | 1 | 30124 |
| PXD014877 | *Clostridium saccharolyticum* | Q Exactive HF-X | Trypsin | 1 | 32736 |
| PXD014877 | *Collinsella Aerofaciens* | Q Exactive HF-X | Trypsin | 1 | 28061 |
| PXD014877 | *Danio rerio* | Q Exactive HF-X | Trypsin | 7 | 340169 |
| PXD014877 | *Eggerthella Lenta* | Q Exactive HF-X | Trypsin | 1 | 11032 |
| PXD014877 | *Fusarium oxysporum* | Q Exactive HF-X | Trypsin | 11 | 185361 |
| PXD014877 | *Fusobacterium Nucleatum Subsp Nucleatum* | Q Exactive HF-X | Trypsin | 1 | 26424 |
| PXD014877 | *Gallus Gallus* | Q Exactive HF-X | Trypsin | 8 | 294035 |
| PXD014877 | *Gossypium hirsutum* | Q Exactive HF-X | Trypsin | 8 | 248653 |
| PXD014877 | *Halobacterium sp* | Q Exactive HF-X | Trypsin | 1 | 31395 |
| PXD014877 | *Lactobacillus Paracasei* | Q Exactive HF-X | Trypsin | 1 | 340169 |
| PXD014877 | *Mus musculus* | Q Exactive HF-X | Trypsin | 7 | 267991 |
| PXD014877 | *Mycoplasmen* | Q Exactive HF-X | Trypsin | 1 | 3608 |
| PXD014877 | *Neurospora Crassa* | Q Exactive HF-X | Trypsin | 7 | 78622 |
| PXD014877 | *Odoribacter splanchnicus* | Q Exactive HF-X | Trypsin | 1 | 40367 |
| PXD014877 | *Parabacteroides Distasonis* | Q Exactive HF-X | Trypsin | 1 | 34529 |
| PXD014877 | *Parabacteroides Merdae* | Q Exactive HF-X | Trypsin | 1 | 39236 |
| PXD014877 | *Phaeodactylum tricornutum* | Q Exactive HF-X | Trypsin | 7 | 126526 |
| PXD014877 | *Prevotella Copri* | Q Exactive HF-X | Trypsin | 1 | 31720 |
| PXD014877 | *Roseburia Intestinalis* | Q Exactive HF-X | Trypsin | 1 | 17139 |
| PXD014877 | *Ruminococcus Bromii* | Q Exactive HF-X | Trypsin | 1 | 20232 |
| PXD014877 | *Ruminococcus Gnavus* | Q Exactive HF-X | Trypsin | 1 | 27689 |
| PXD014877 | *Streptococcus Parasanguinis* | Q Exactive HF-X | Trypsin | 1 | 26601 |
| PXD014877 | *Sulfolobus solfataricus* | Q Exactive HF-X | Trypsin | 1 | 1926 |
| PXD014877 | *Symbiodinium microadriaticum* | Q Exactive HF-X | Trypsin | 5 | 4310 |
| PXD014877 | *Triticum aestivum* | Q Exactive HF-X | Trypsin | 7 | 265766 |
| PXD014877 | *Veillonella parvula* | Q Exactive HF-X | Trypsin | 1 | 30734 |
| PXD014877 | *Vitris vinifera* | Q Exactive HF-X | Trypsin | 7 | 203696 |
| PXD014877 | *Akkermansia muciniphilia* | Q Exactive HF-X | Trypsin | 1 | 22013 |
| PXD014877 | *Bacteroides Thetaiotaomicron* | Q Exactive HF-X | Trypsin | 1 | 50460 |
| PXD014877 | *Drosophila* | Q Exactive HF-X | Trypsin | 8 | 217125 |
| PXD014877 | *Ruminococcus Torques* | Q Exactive HF-X | Trypsin | 1 | 20811 |
| PXD016986 | *Riftia pachyptila* | LTQ Orbitrap Velos | Trypsin | 349 | 1973461 |
| PXD018830 | *Homo sapiens* | Q Exactive Plus | Trypsin | 276 | 2527193 |
| PXD019483 | *hCOV* | Q Exactive HF-X | Trypsin | 1 | 151 |
| PXD019483 | *HSV* | Q Exactive HF-X | Trypsin | 1 | 3288 |
| PXD019483 | *LACV* | Q Exactive HF-X | Trypsin | 1 | 2339 |
| PXD019483 | *RVFV* | Q Exactive HF-X | Trypsin | 2 | 1429 |
| PXD019483 | *VSV* | Q Exactive HF-X | Trypsin | 1 | 819 |
| PXD019483 | *ZIKV* | Q Exactive HF-X | Trypsin | 2 | 835 |
| PXD019483 | *Homo sapiens* | Q Exactive HF-X | Trypsin | 111 | 4001686 |
| PXD019643 | *Homo sapiens* | Orbitrap Fusion Lumos | Trypsin | 719 | 4290071 |
| PXD020192 | *Homo sapiens* | Q Exactive HF-X | Trypsin | 91 | 7649835 |
| PXD021013 | *Homo sapiens* | Orbitrap Fusion Lumos | Trypsin | 996 | 16314438 |
| PXD022469 | *Homo sapiens* | Q Exactive Plus | Trypsin | 383 | 696434 |
| PXD024364 | *Homo sapiens* | Orbitrap Fusion | Lys-C | 30 | 509465 |
| PXD024364 | *Homo sapiens* | Orbitrap Fusion | Trypsin | 281 | 7120784 |
| PXD028735 | *Escherichia coli* | Q Exactive HF-X | Trypsin | 39 | 99467 |
| PXD028735 | *Homo sapiens* | Q Exactive HF-X | Trypsin | 45 | 1511325 |
| PXD030348 | *Mus musculus* | Orbitrap Fusion | Trypsin | 284 | 6483317 |
| PXD030983 | *Mus musculus* | Q Exactive HF | Trypsin | 2137 | 30714490 |
| PXD040266 | *Mus musculus* | timsTOF | Trypsin | 48 | 896379 |
| PXD010429 | *Homo sapiens* | Q Exactive Plus | Trypsin | 696 | 2578074 |
| PXD005478 | *Mycobacterium tuberculosis* | Q Exactive HF | Trypsin | 582 | 2013520 |
| PXD002899 | *Xenopus laevis* | Q Exactive HF | Trypsin | 357 | 3498190 |
| PXD004352 | *Homo sapiens* | Q Exactive Plus | Trypsin | 455 | 1208405 |
| PXD025555 | *Homo sapiens* | Orbitrap Fusion Lumos | Trypsin | 797 | 9620961 |
| PXD005526 | *Mus musculus* | Q Exactive | Trypsin | 395 | 13527205 |
| PXD021398 | *Homo sapiens* | Q Exactive | Trypsin | 566 | 4305190 |
| PXD034656 | *Saccharomyces cerevisia* | Q Exactive HF | Trypsin | 639 | 3264025 |
| PXD005252 | *Homo sapiens* | Q Exactive HF | Trypsin | 87 | 765297 |
| PXD041268 | *Homo sapiens* | Orbitrap Eclipse | Trypsin | 347 | 11322589 |
| PDC000219 | *Homo sapiens* | Orbitrap Fusion Lumos | Trypsin | 1288 | 25716227 |
| PXD046290 | *Balneola vulgaris, Deinococcus proteolyticus mrp, Homo sapiens* | Orbitrap Astral | Trypsin | 12 | 1364149 |
| PXD062529 | *Homo sapiens* | Q Exactive HF | Trypsin | 3833 | 30248716 |
| PDC000127 | *Homo sapiens* | Orbitrap Fusion Lumos | Trypsin | 575 | 7515500 |
| PDC000220 | *Homo sapiens* | Orbitrap Fusion Lumos | Trypsin | 504 | 1871613 |
| PXD046320 | *Homo sapiens* | Orbitrap Astral | Trypsin | 6 | 142938 |
| PXD054415 | Metaproteomics*:* *Escherichia coli K12, Thermus Thermophilus, et al* | Orbitrap Fusion | Trypsin | 36 | 1407405 |
| PXD059114 | Metaproteomics*:* *Nitrosomonas europaeae,* *Nitrosomonas ureae, et al* | timsTOF | Trypsin | 357 | 6807463 |
| IPX0010296000 | *Homo sapiens* | Orbitrap Astral | Trypsin | 515 | 50587097 |
| HuiYanMouse | *Mus musculus* | Orbitrap Astral | Trypsin | 466 | 33944576 |
| PXD054790 | *Homo sapiens* | Q Exactive HF | Trypsin | 1201 | 64804704 |

Supplementary Table 2: The core parquet structure of π-MSNet PSM results

| column_name | data_type | description |
| --- | --- | --- |
| sequence | string | The peptide's sequence (with no modifications) |
| peptidoform | string | Peptide sequence with modifications, see more in the documentation |
| modifications | list<item: struct<name: string, accession: string, positions: list<item: struct<position: string, scores: list<item: struct<score_name: string, score_value: double, higher_better: bool>>>>>> | List of modifications with details on position and position-specific scores |
| precursor_charge | int32 | Precursor charge |
| posterior_error_probability | double | Posterior error probability for a given peptide or PSM. |
| is_decoy | int32 | Decoy indicator, 1 if the PSM is a decoy, 0 target |
| cal_mass_to_charge | float | Theoretical peptide mass-to-charge ratio based on identified sequence and modifications |
| exp_mass_to_charge | float | Experimental peptide mass-to-charge ratio of identified peptide |
| reference_file_name | string | The reference file name that contains the feature |
| cv_params | list<item: struct<cv_name: string, cv_value: string>> | Optional list of CV parameters for additional metadata |
| scan | string | Scan number of the spectrum |
| retention_time | float | MS2 scan's precursor retention time (in seconds) |
| ion_mobility | float | Ion mobility value for the precursor ion |
| mz_array | list<item: float> | Array of m/z values for the spectrum used for the peptide spectrum match |
| intensity_array | list<item: float> | Array of intensity values for the spectrum used for the peptide spectrum match |
| charge_array | list<item: int32> | Array of fragment ion charge values for the spectrum used for the peptide spectrum match |
| ion_type_array | list<item: string> | Array of fragment ion type annotations (e.g., b, y, a) for the spectrum used for the peptide spectrum match |

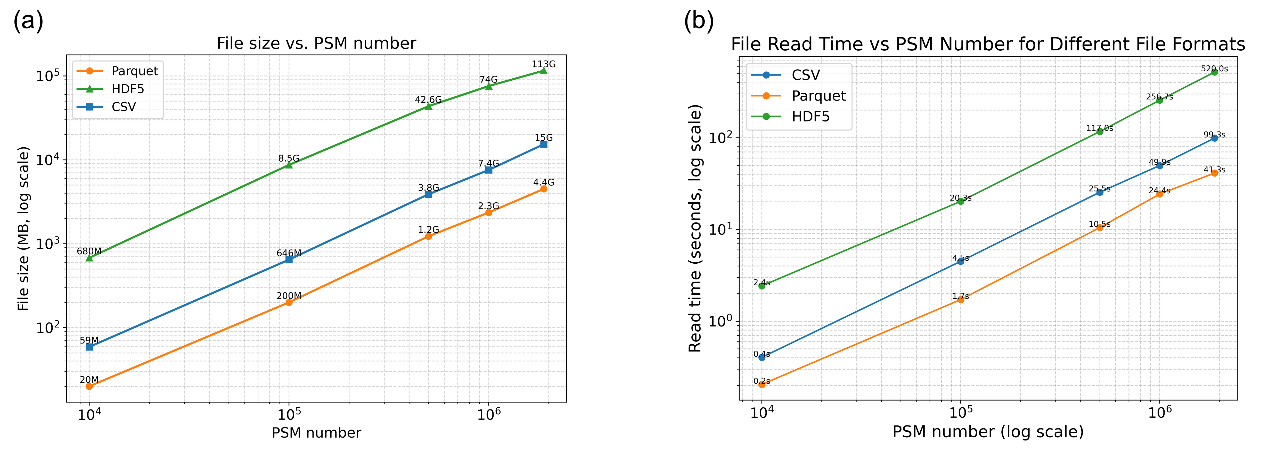

Supplementary Figure 2: Comparison of storage efficiency and read performance across file formats**.** (a) File size (MB, log scale) as a function of peptide-spectrum match (PSM) number for three file formats (CSV, Parquet, and HDF5). Parquet consistently achieves the smallest file size, followed by CSV, while HDF5 exhibits the largest storage overhead. (b) File read time (seconds, log scale) versus PSM number for the same formats. Parquet demonstrates the fastest read performance across all dataset sizes, while HDF5 shows substantially slower read times, particularly at larger scales. The benchmark data were randomly sampled from the π-HuB Orbitrap Exploris 480 dataset^4^.

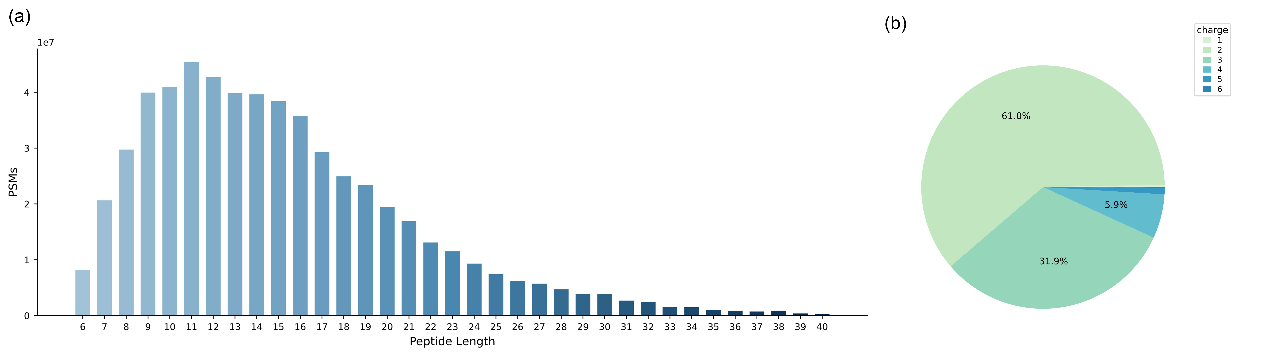

Supplementary Figure 3: Distribution of the π-MSNet dataset. (a) The number of peptide-spectrum matches (PSMs) across different peptide sequence lengths. (b) Precursor charge state distribution in π-MSNet. The majority of spectra correspond to charge states 2 and 3, whereas higher charge states are less frequent.

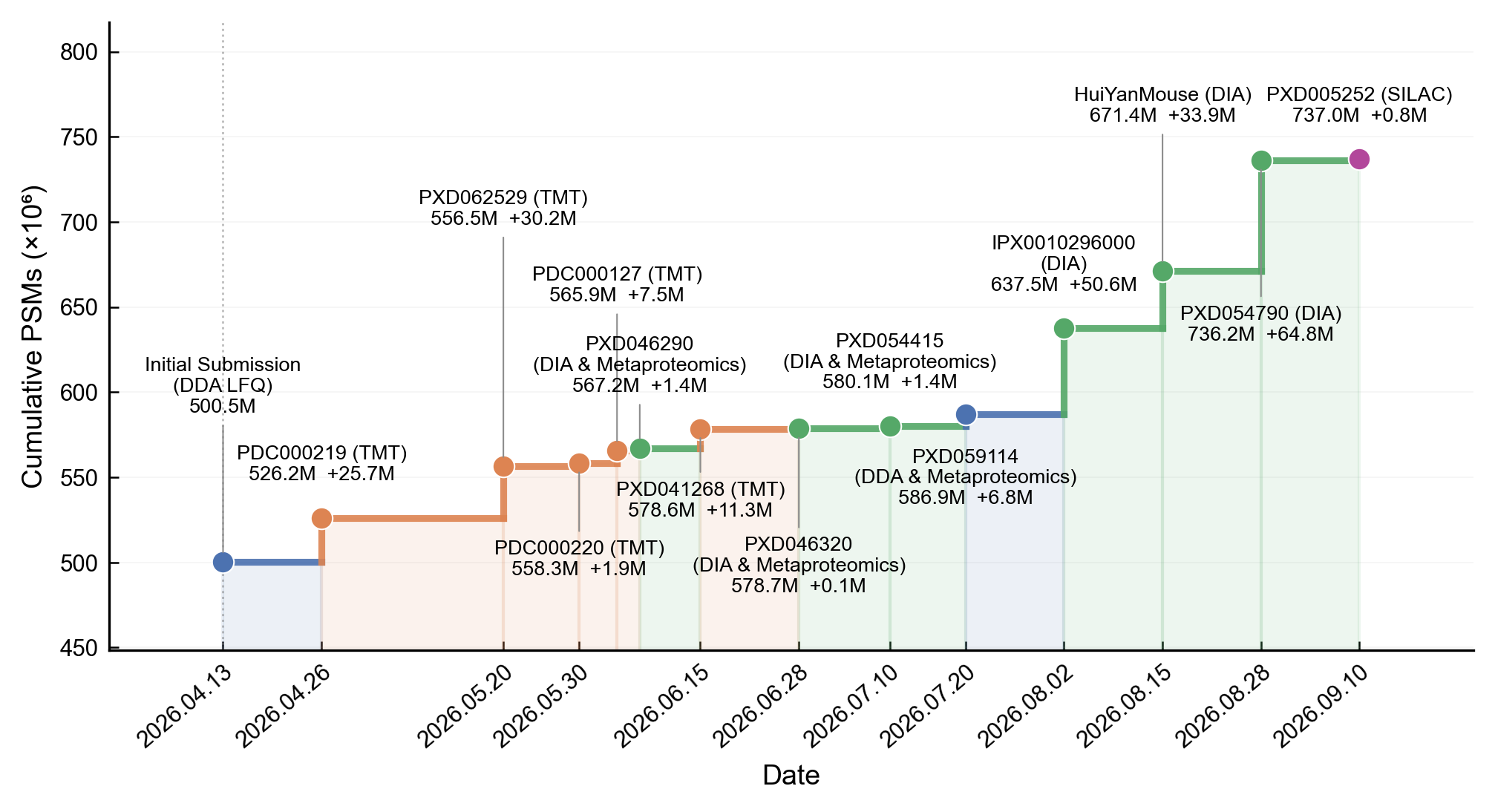

Supplementary Figure 4: Cumulative increase in released PSMs since manuscript submission. The step curve traces total spectral count from the initial 500.5 million PSMs (DDA LFQ, 2026.04.13) to the final 737 million PSMs (2026.09.10), incorporating 13 incremental public deposits. Point colors encode dataset type: blue — DDA LFQ and DDA metaproteomics, orange — TMT, green — DIA metaproteomics, purple — SILAC. The vertical dotted line indicates the submission date. DDA: data-dependent acquisition, DIA: data-independent acquisition, LFQ: label-free quantification, TMT: tandem mass tag, SILAC: Stable isotope labeling by amino acids in cell culture.

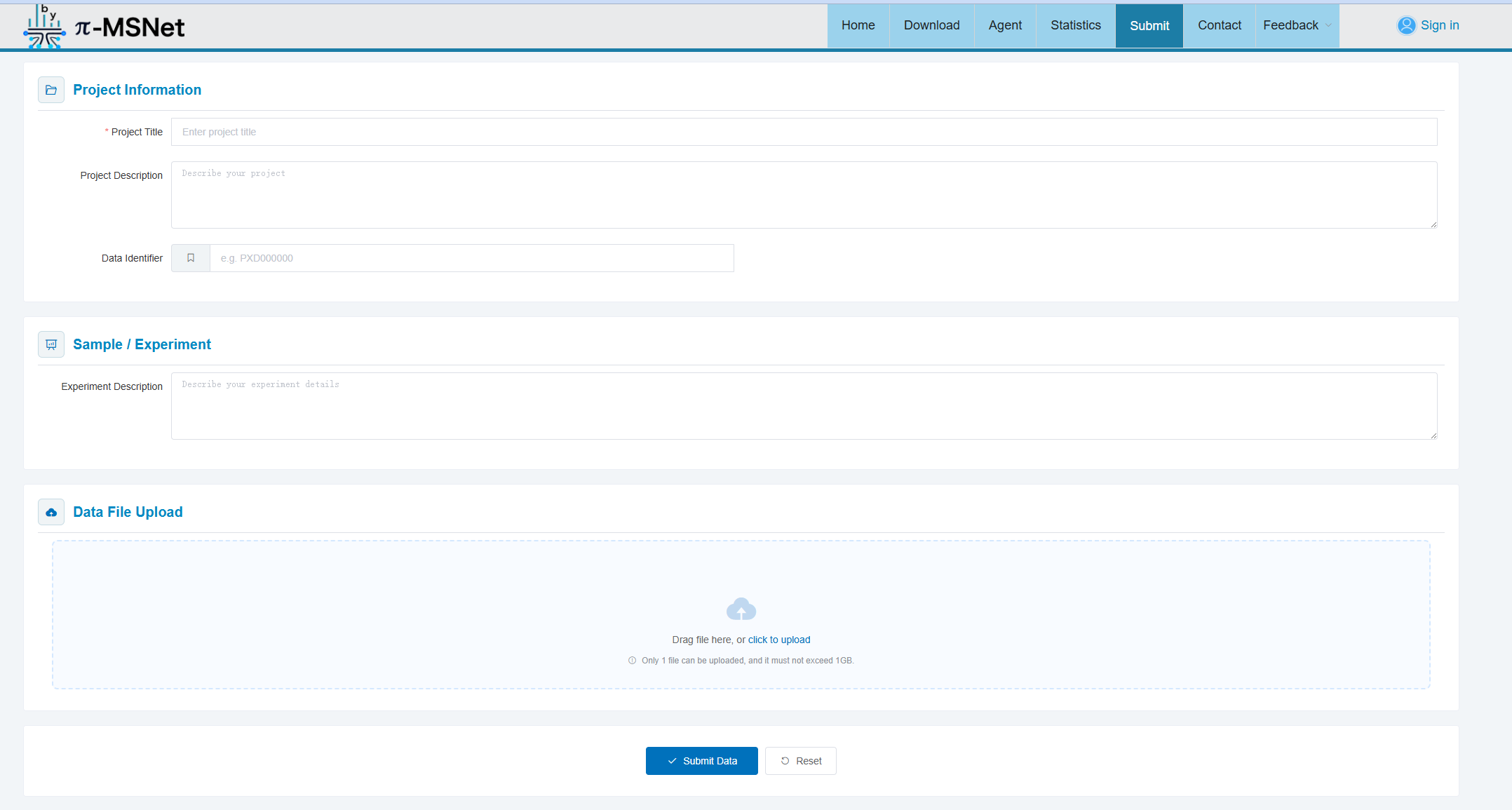

Supplementary Figure 5: Standardized deposition portal via web interface.

Supplementary Note 2: MS2 prediction model training and benchmark.

We evaluated MS2 prediction models across heterogeneous π-MSNet datasets and further examined scaling behavior with increasing training data, training steps, and model size. The datasets used for model training and benchmarking are summarized in Supplementary Table 3, with additional rescoring results shown in Supplementary Figure 6.

Supplementary Table 3. π-MSNet Datasets Used in Evaluation of Fragmentation Intensity Profiles**.** Heterogeneous MS data are used for benchmarking, including samples that have been using trypsin and alternative proteases; samples originating from various species, including human, animals, plants and prokaryotes; data that acquired using different MS platform from multiple instrument vendors.

|  | **Dataset** | **Orbitrap Elite** | **Orbitrap Velos** | **Q Exactive HF** | **Q Exactive HF-X** | **Exploris480** | **Orbitrap Fusion Lumos** | **Q Exactive** | **Q Exactive Plus** | **Orbitrap Fusion** | **Human**  **Animal**  **Plant**  **Prokaryote**  **Other** | | | | | **NCE** | **#PSM** | **#Precursor** |
| --- | --- | --- | --- | --- | --- | --- | --- | --- | --- | --- | --- | --- | --- | --- | --- | --- | --- | --- |
| **MS2 train datasets** | PXD004732 |  |  |  |  |  |  |  |  |  |  |  |  |  |  | 30&25 | 7930592 | 278199 |
|  | PXD010595 |  |  |  |  |  |  |  |  |  |  |  |  |  |  | 35&30&25&23&20 | 5459100 | 154339 |
|  | Hub E480 Cohort |  |  |  |  |  |  |  |  |  |  |  |  |  |  | 30&28 | 501476 | 44665 |
|  | PXD012131 |  |  |  |  |  |  |  |  |  |  |  |  |  |  | 30&25 | 541775 | 65902 |
|  | PXD014877 Drosophila& Triticum_aestivum& Gossypium |  |  |  |  |  |  |  |  |  |  |  |  |  |  | 27 | 18240 | 9255 |
|  | PXD006675 |  |  |  |  |  |  |  |  |  |  |  |  |  |  | 27 | 550266 | 75394 |
|  | PXD010899 |  |  |  |  |  |  |  |  |  |  |  |  |  |  | 30 | 135884 | 24471 |
|  | PXD030983 |  |  |  |  |  |  |  |  |  |  |  |  |  |  | 28&25 | 1953642 | 156996 |
|  | PXD004242 |  |  |  |  |  |  |  |  |  |  |  |  |  |  | 27 | 72424 | 8050 |
|  | PXD008722 |  |  |  |  |  |  |  |  |  |  |  |  |  |  | 28 | 741554 | 111819 |
|  | PXD008840 |  |  |  |  |  |  |  |  |  |  |  |  |  |  | 27 | 358674 | 89866 |
|  | PXD028735-Ecoli |  |  |  |  |  |  |  |  |  |  |  |  |  |  | 27 | 60672 | 8647 |
|  | PXD004452 Gluc |  |  |  |  |  |  |  |  |  |  |  |  |  |  | 28 | 117602 | 82098 |
|  | PXD024364 lysc |  |  |  |  |  |  |  |  |  |  |  |  |  |  | 30 | 376304 | 117952 |
|  | PXD021013 |  |  |  |  |  |  |  |  |  |  |  |  |  |  | 35&  30&  25&23&20 | 7603021 | 489329 |
|  | PXD013868 |  |  |  |  |  |  |  |  |  |  |  |  |  |  | 25 | 1136413 | 121309 |
|  | PXD019643 |  |  |  |  |  |  |  |  |  |  |  |  |  |  | 30 | 2448050 | 143961 |
|  | PXD012636 |  |  |  |  |  |  |  |  |  |  |  |  |  |  | 28 | 114739 | 5438 |
|  | PXD010154 |  |  |  |  |  |  |  |  |  |  |  |  |  |  | 25 | 1454748 | 275145 |
|  | IPX0001804001 |  |  |  |  |  |  |  |  |  |  |  |  |  |  | 32 | 105490 | 19264 |
|  | IPX0002031000 |  |  |  |  |  |  |  |  |  |  |  |  |  |  | 27 | 191849 | 30773 |
|  | PXD002767 |  |  |  |  |  |  |  |  |  |  |  |  |  |  | 30 | 2419261 | 77271 |
|  | PXD062529 |  |  |  |  |  |  |  |  |  |  |  |  |  |  | 30 | 4167793 | 243076 |
| **MS2 test datasets** | PXD012636 |  |  |  |  |  |  |  |  |  |  |  |  |  |  | 28 | 63262 | 13604 |
|  | IPX0004073001 |  |  |  |  |  |  |  |  |  |  |  |  |  |  | 30 | 411712 | 5518 |
|  | PXD019483 |  |  |  |  |  |  |  |  |  |  |  |  |  |  | 27 | 1612324 | 163989 |
|  | PXD014877 |  |  |  |  |  |  |  |  |  |  |  |  |  |  | 27 | 219766 | 102217 |
|  | PXD009737 Gluc/LysC |  |  |  |  |  |  |  |  |  |  |  |  |  |  | 27 | 328597 | 138774 |
|  | PXD000561 |  |  |  |  |  |  |  |  |  |  |  |  |  |  | 32 | 1382583 | 270118 |
|  | HuiYan |  |  |  |  |  |  |  |  |  |  |  |  |  |  | 30&28 | 856541 | 115080 |
|  | PXD062529 |  |  |  |  |  |  |  |  |  |  |  |  |  |  | 30 | 168196 | 23013 |

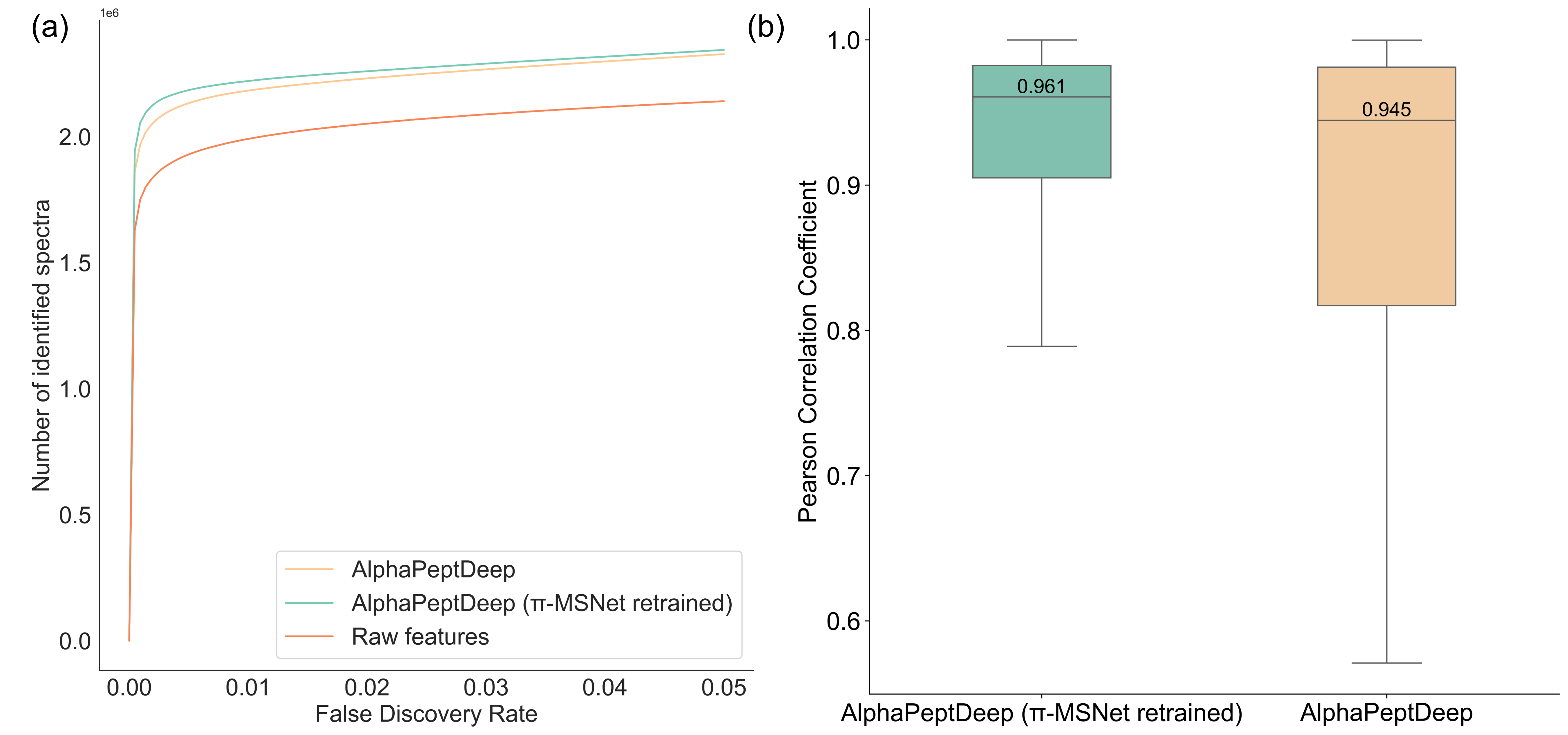

Supplementary Figure 6: Performance comparison of PSM rescoring on the dataset PXD010154 using the original and π-MSNet-retrained AlphaPeptDeep models. (a) The number of identified PSMs were obtained before and after prediction-based rescoring with Percolator^5^ using raw AlphaPeptDeep^6^ model weights and π-MSNet model weights for various q-values. (b) The distribution of Pearson Correlation Coefficient (PCC) between observed and predicted spectrum using raw AlphaPeptDeep model weights and π-MSNet model weights.

Supplementary Note 3: Retention time prediction model training and benchmark.

We evaluated RT prediction models using two data-splitting strategies and further examined their scaling behavior with increasing training data. Additional scaling-law results and the characteristics of the test groups are shown in Supplementary Figures 7 and 8.

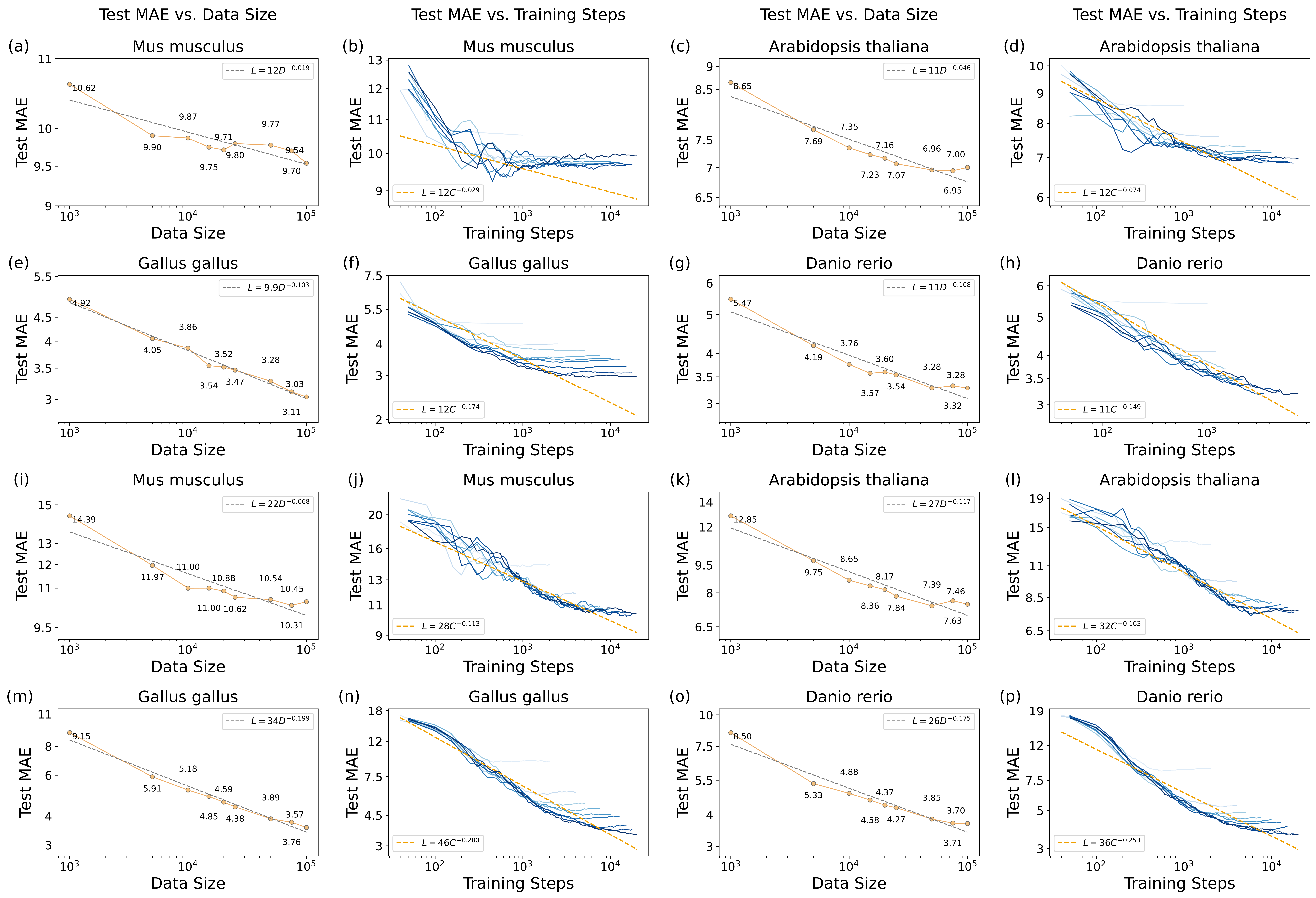

Supplementary Figure 7: Scaling-law analyses for the remaining held-out species. (a-h) show AutoRT^7^ scaling-law results for held-out *Mus musculus* (a,b), *Arabidopsis thaliana* (c,d), *Gallus gallus* (e,f), and *Danio rerio* (g,h). (i-p) show the corresponding DeepLC^8^ results for *Mus musculus* (i,j), *Arabidopsis thaliana* (k,l), *Gallus gallus* (m,n), and *Danio rerio* (o,p). For each species, the left plot shows converged MAE as a function of training-set size, whereas the right plot shows test MAE across training steps for different training-set sizes, with darker loss curves indicating larger training-set sizes. Models were trained from scratch using nested training subsets sampled from the corresponding non-held-out-species pool and evaluated on a randomly sampled 1,000-peptide test set from the held-out species. Dashed lines indicate log-log power-law fits.

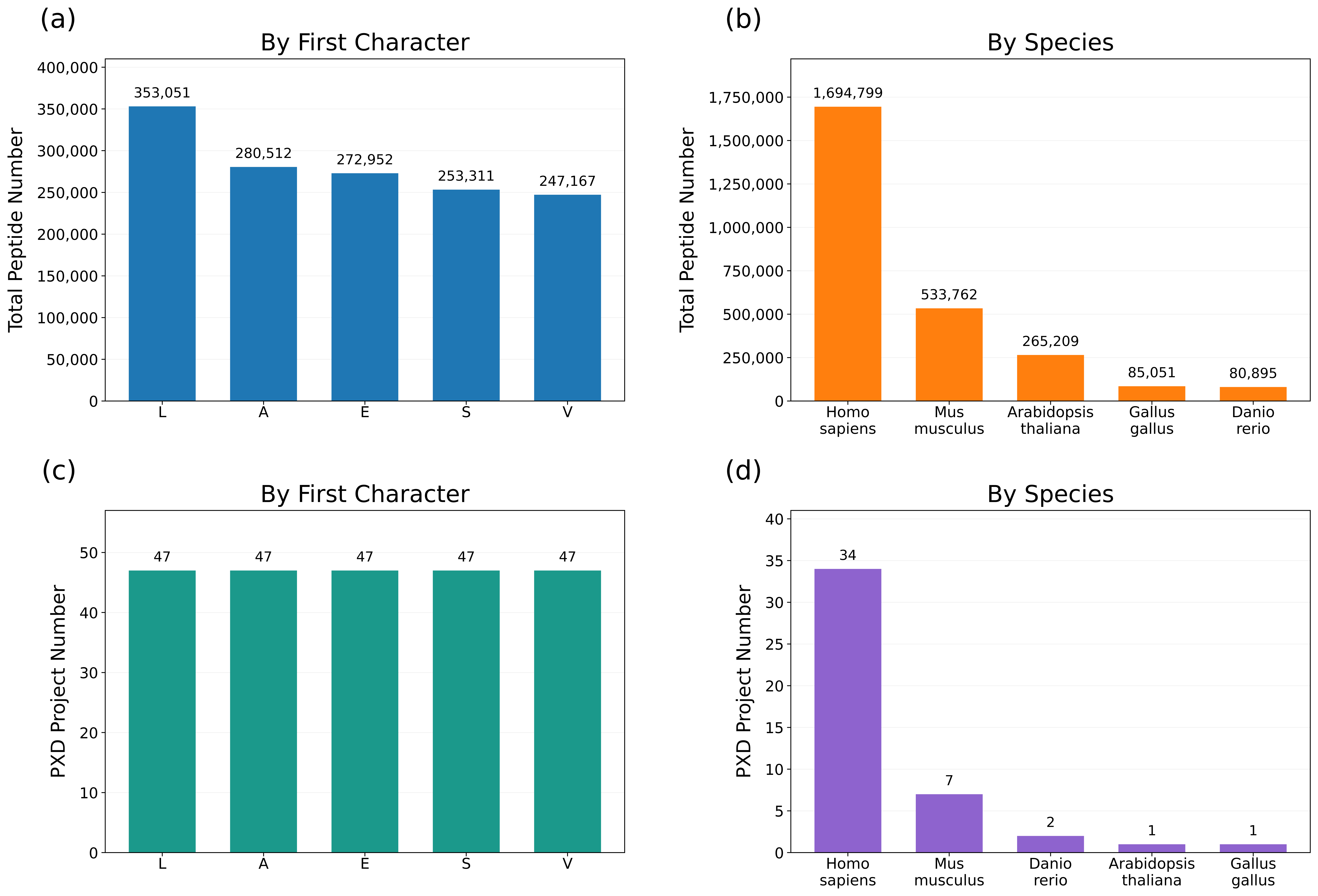

Supplementary Figure 8: Numbers of peptides and source ProteomeXchange (PXD) projects in the test groups defined by the two data-splitting strategies used for RT prediction. (a) and (b) show the peptide counts for the top five N-terminal amino acid groups and the top five species groups, respectively. In the N-terminal amino acid-based splitting strategy, peptides beginning with a specific amino acid were assigned to the test set, whereas all remaining peptides were used for training. In the species-based splitting strategy, peptides from a specific species were assigned to the test set, whereas peptides from the remaining species were used for training. (c) and (d) show the corresponding numbers of distinct source PXD projects.

Supplementary Note 4: Scaling laws for collisional cross sections prediction model.

The PXD019086 dataset was used to construct the training and test data for CCS prediction. For precursors with repeated measurements, the CCS values were averaged to obtain a single value. A total of 465,356 unique precursors were used as the training set, and an additional 168,570 non-overlapping precursors were used as an independent test set. For data-scaling experiments, the model architecture was fixed and models were trained from random initialization using nested training subsets of 500, 1,000, 2,500, 5,600, 10,000, 25,000, 50,000, 100,000, 200,000, 280,000, 400,000 and 465,356 precursors. Three random seeds were used for each training-set size. For compute-scaling experiments, the model architecture was fixed and learning curves were evaluated across different training steps. For parameter-scaling experiments, the training-set size was fixed at the maximum of 465,356 precursors, while the hidden dimension of the model was varied to generate models with different numbers of trainable parameters. Each model was trained from random initialization using three random seeds.

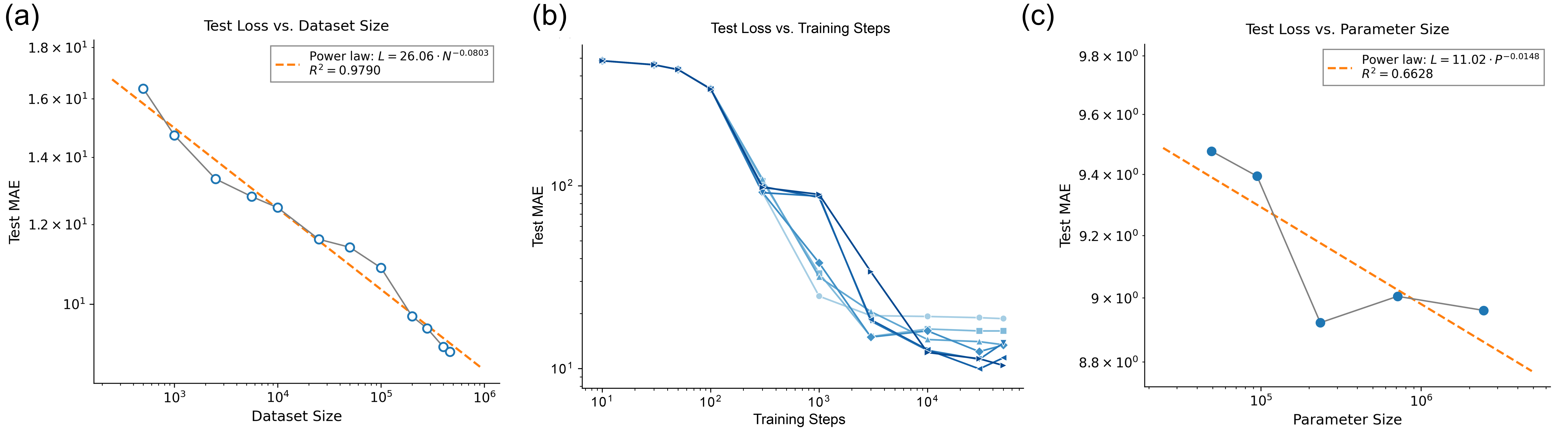

Supplementary Figure 9: Scaling law validation in CCS prediction. (a) Dataset size, (b) computation steps, and (c) model size used for training the AlphaPeptDeep CCS model^9^. Test losses represent the mean across three random seeds. In (b), lines represent different training dataset sizes, with darker colours indicating larger datasets.

Supplementary Note 5: Architecture of the π-MSNet agent.

The π-MSNet agent provides a natural-language interface to proteomics analysis workflows (Supplementary Figure 10). Users submit a request with the relevant input, such as a pasted precursor list for MS2 spectrum prediction, an MGF file for de novo peptide sequencing or a peptide list for retention time prediction. Built on the OpenAI Agents SDK with a deepseek-v4-flash reasoning engine, the agent identifies the user’s intent, selects the appropriate task-specific skill and submits specific jobs to the server. The supported outputs include predicted MS2 spectra, peptide sequences, retention times and DIA libraries and its visualization artifacts.

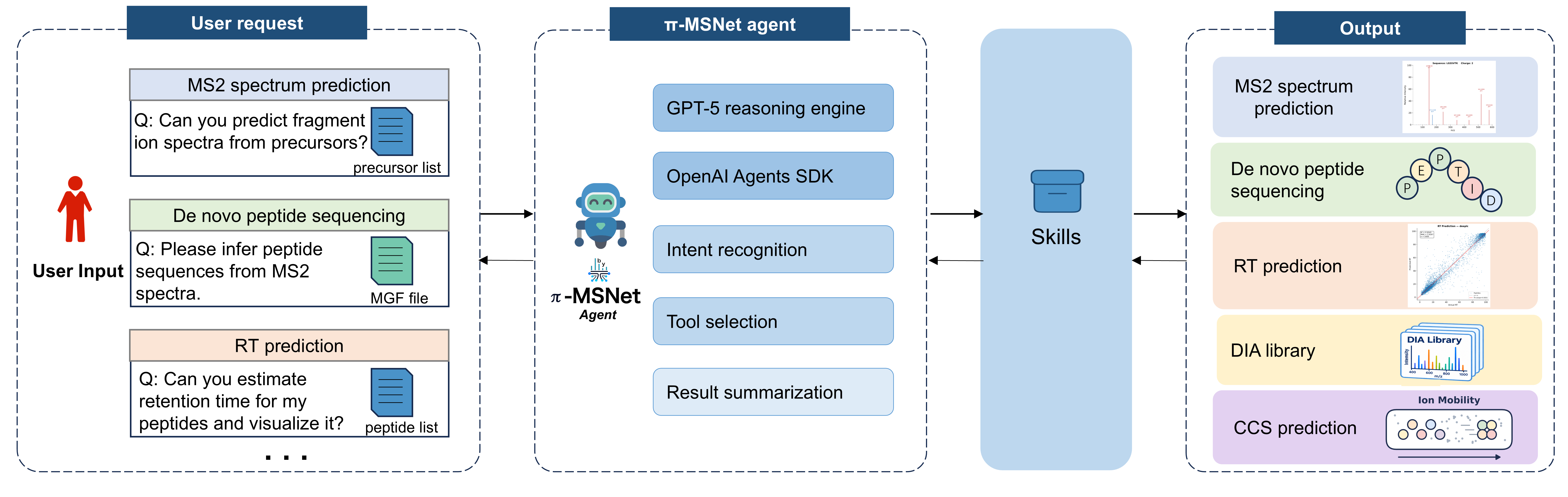

Supplementary Figure 10: Schematic illustration of the π-MSNet agent framework. Users can interact with the agent via natural language to perform multiple proteomics tasks, including MS2 spectrum prediction, de novo peptide sequencing, and retention time (RT) prediction, along with corresponding visualization outputs.

**Supplementary References**

1. Dai, C. et al. quantms: a cloud-based pipeline for quantitative proteomics enables the reanalysis of public proteomics data. Nat Methods 21, 1603–1607 (2024).
2. Yue, Q. et al. quantmsdiann: a scalable SDRF-driven DIA-NN workflow for reanalysis of single-cell, spatial, and bulk proteomics datasets. Research Square (2026).
3. Demichev, V., Messner, C. B., Vernardis, S. I., Lilley, K. S. & Ralser, M. DIA-NN: neural networks and interference correction enable deep proteome coverage in high throughput. Nat Methods 17, 41–44 (2020).
4. He, F. et al. π-HuB: the proteomic navigator of the human body. Nature 636, 322–331 (2024).
5. Käll, L., Canterbury, J. D., Weston, J., Noble, W. S. & MacCoss, M. J. Semi-supervised learning for peptide identification from shotgun proteomics datasets. Nat Methods 4, 923–925 (2007).
6. Zeng, W.-F. et al. AlphaPeptDeep: a modular deep learning framework to predict peptide properties for proteomics. Nat Commun 13, 7238 (2022).
7. Maboudi Afkham, H., Qiu, X., The, M. & Käll, L. Uncertainty estimation of predictions of peptides’ chromatographic retention times in shotgun proteomics. Bioinformatics 33, 508–513 (2017).
8. Bouwmeester, R., Gabriels, R., Hulstaert, N., Martens, L. & Degroeve, S. DeepLC can predict retention times for peptides that carry as-yet unseen modifications. Nat Methods 18, 1363–1369 (2021).
9. Meier, F. et al. Deep learning the collisional cross sections of the peptide universe from a million experimental values. Nat Commun 12, 1185 (2021).
